# *In Vivo* Efficacy of ABHD17 Inhibition Validates the Palmitoylation Cycle as a Therapeutic Target in *NRAS*-Mutant Cancer

**DOI:** 10.1101/2025.03.20.644389

**Authors:** Matthew Decker, Benjamin J. Huang, Timothy Ware, Christopher Boone, Michelle Tang, Julia Ybarra, Matthew J. Sale, Rony A. François, Aishwarya C. Ballapuram, Katrine A. Taran, Esteban Tarazona-Guzman, Pan-Yu Chen, Marcos Armendáriz, Camille J. Leung, Max Harris, Karensa Tjoa, Henry Hongo, Sydney Abelson, Jose Rivera, Johanna Virta, Nhi Ngo, Dylan M. Herbst, Radu M. Suciu, Carlos Guijas, Kimia Sedighi, Taylor Andalis, Elysia Roche, Boer Xie, Yunlong Liu, Catherine C. Smith, Michelle Arkin, Elliot Stieglitz, Frank McCormick, Micah J. Niphakis, Benjamin F. Cravatt, Kevin Shannon

**Author notes:** These authors contributed equally (Matthew Decker, Benjamin J. Huang, Timothy Ware).

## Abstract

Ras proteins are functionally dependent on one or more lipid modifications.^1,2^ The dynamic palmitoylation of N-Ras by DHHC palmitoyl acyltransferases and depalmitoylation by ABHD17 serine hydrolases is essential for the growth of *NRAS*-mutant acute myeloid leukemia (AML) cells.^3–6^ Here we show that ABD778, an *in vivo*-active ABHD17 inhibitor, selectively reduces the growth of *NRAS*-mutant AML and melanoma cell lines and is synergistic with the MEK inhibitor PD0325901 (PD901; mirdametinib). Mechanistically, ABD778 and PD901 induce deep and durable suppression of mitogen activated protein kinase (MAPK) pathway activation. Co-treatment extended the survival of mice transplanted with *Nras^G12D^* AMLs, which acquired by-pass mutations at relapse that conferred drug resistance and restored MAPK activation. ABD778 augmented the anti-leukemia activity of PI3 kinase, pan-Ras tri-complex, and FLT3 inhibitors, and restored gilteritinib sensitivity in a patient-derived xenograft model of FLT3 inhibitor resistance. These studies validate the palmitoylation cycle as a therapeutic target in *NRAS*-mutant cancers.

## INTRODUCTION

Somatic *NRAS* mutations occur in 3-5% of adult and pediatric cancers and are most prevalent in hematologic malignancies, melanoma, and thyroid cancer.^7,8^ In addition, *NRAS* mutations are key drivers of *de novo* and adaptive resistance to BCL-2, IDH1/2, and FLT3 inhibitors in acute myeloid leukemia (AML) and in melanomas that relapse after Raf kinase inhibitor treatment.^9–12^ There are no approved targeted therapies for *NRAS*-mutant cancers.

Sequential farnesylation and palmitoylation of the N-Ras C-terminal hypervariable region (HVR) mediates stable association with cell membranes and is required for efficient N-Ras activation and downstream signaling.^1^ Whereas farnesylation is irreversible, the N-Ras palmitoylation/depalmitoylation cycle is highly dynamic.^13^ The depalmitoylation reaction maintains a kinetic trap in the Golgi apparatus where addition of the palmitoyl group significantly increases lipophilic affinity of N-Ras and locks it in place.^14^ Once it is trapped in the Golgi via palmitoylation, N-Ras undergoes anterograde trafficking to the plasma membrane in a complex with VPS35.^15^ Inhibiting either the “on” or the “off” reaction impairs downstream signaling.^16^

The essential role of farnesylation in Ras transformation stimulated drug discovery programs in the 1990s that generated potent and selective farnesyl transferase inhibitors.^17^ Unfortunately, these compounds proved ineffective in most *RAS-*mutant cancers due to by-pass geranylgeranylation of the C-terminal CaaX motifs of N-Ras and K-Ras.^17–20^ After this disappointing experience, the pharmaceutical industry largely abandoned efforts to develop new approaches for inhibiting the lipid modification of N-Ras and K-Ras.

The palmitoylation cycle has theoretical advantages over farnesylation/geranylgeranylation as a therapeutic target in *NRAS-*mutant cancers. No compensatory lipid modification for N-Ras palmitoylation has yet been identified. In addition, while the palmitoylation cycle is required for proper localization and activity of N-Ras, K-Ras4a, and H-Ras, the K-Ras4b HVR has a lipophilic poly-lysine domain that enhances membrane affinity, which obviates the requirement for a second lipid modification. In principle, this could represent a “biologic therapeutic index” by allowing non-tumor tissues to tolerate chemical inhibition of the N-Ras palmitoylation cycle by signaling through K-Ras4b. Finally, we and others showed that introducing a cysteine-to-serine mutation (C181S) into the N-Ras HVR abrogates palmitoylation, inhibits the growth of *NRAS*-mutant AML cell lines in a genotype-specific manner, and suppresses myeloid transformation by oncogenic *NRAS in vivo.*^3,5,6,21^ While these data provide a rationale for developing chemical inhibitors of the DHHC enzymes that palmitoylate N-Ras, DHHC9 is the only known N-Ras PAT and multiple lines of evidence support a physiologic role for additional family members.^6,22,23^

The ABHD17 serine hydrolases are the principal N-Ras depalmitoylases.^4,5^ We previously developed ABD957, a pan-ABHD17 inhibitor that selectively suppressed MAPK signaling and reduced the growth of cultured *NRAS-*mutant AML cell lines.^5^ ABD957 also showed synergistic anti-leukemia activity with the allosteric MEK inhibitor PD0325901 (PD901; mirdametinib), which is approved by the US Food and Drug Administration (FDA) for the treatment of plexiform neurofibroma in patients with neurofibromatosis, type 1. The promising *in vitro* activity of ABD957 inspired efforts to develop and characterize an ABHD17 inhibitor suitable for *in vivo* studies that culminated in the discovery of the potent, selective and *in vivo*-active ABHD17 inhibitor ABD778.

Here we show that co-treatment with ABD778 and PD901 selectively inhibits the growth of AML cell lines and primary juvenile myelomonocytic leukemia (JMML) cells harboring oncogenic *NRAS* mutations, markedly extends the survival of recipient mice transplanted with *Nras-*mutant AMLs, and drives the emergence of adaptive resistance *in vivo.* Biochemical analysis revealed prolonged MAPK pathway inhibition in response to ABD778 and PD901 and support a model whereby ABD778 decreases Raf activation by N-Ras oncoproteins and blunts feedback reactivation of MAPK signaling in response to MEK inhibition. Co-treatment with ABD778 and PD901 caused similar synergistic growth repression and feedback inhibition in *NRAS*-mutant melanoma cell lines. Additionally, ABD778 exhibited synergy with the class 1 PI3 kinase inhibitor pictilisib^24^, the K/N-Ras^G12C^ inhibitor sotorasib^25^, the Ras tri-complex inhibitor RMC-6236 (daraxonrasib)^26^ and the FLT3 inhibitor gilteritinib in human AML cells expressing oncogenic N-Ras proteins. Co-treatment with ABD778 and gilteritinib also suppressed the growth of a patient-derived xenograft (PDX) model of adaptive resistance to FLT3 inhibition.

## RESULTS

### Development of a pan-ABHD17 inhibitor with *in vivo* activity

Previously, we reported the development of the covalent inhibitor ABD957 that potently inhibited ABHD17 enzymes in cells with high selectivity over other depalmitoylases, including LYPLA1/2, PPT1 and ABHD10.^5^ While ABD957 provided a useful tool to study ABHD17 enzymes in cellular contexts, this compound was found to have low permeability (Caco-2 P_app_ (A-B) = 0.6 cm^-6^/s) and high efflux (efflux ratio = 16.4) in Caco-2 cells that limited its exposure *in vivo*. We reasoned that reducing the polarity of ABD957 by esterifying its carboxylic acid would improve these properties. The isopropyl ester analog of ABD957, here termed ABD778, exhibited enhanced passive permeability (Caco-2 P_app_ (A-B) = 1.3 cm^-6^/s) and minimal efflux (efflux ratio = 1.45) (**Figure 1A**). We next determined if this modification altered the potency and selectivity of ABD778 in mouse brain proteomes and human ABHD17B-transfected HEK293T cell lysates using gel-based activity-based protein profiling (ABPP) where over 25 native serine hydrolases can be evaluated owing to their reactivity with the active site-directed fluorophosphonate rhodamine (FP-Rh) probe.^27,28^ This analysis showed that ABD778 maintained potency against human and mouse ABHD17 enzymes (IC_50_ = 0.076 and 0.084 μM, respectively) (**Figure 1B**) and demonstrated good ABHD17 specificity across the serine hydrolases detected by gel-ABPP in both mouse (**Figure 1C**) and human (**Figure 1D**) proteomes. Deeper selectivity profiling was then performed across >90 mouse and human serine hydrolases in mouse brain and OCI-AML3 (a human *NRAS*-mutant AML cell line) proteomes using mass spectrometry (MS)-ABPP,^29^ which identified a few additional ABD778-reactive serine hydrolases, including ABHD6 and several carboxyesterases (**Figures S1A, S1B**). Notably, these primary off-targets are common to many covalent serine hydrolase inhibitors^30^ and were accounted for in previous experiments with ABD957 through the use of inactive control compounds,^5^ which revealed that these off-targets do not substantially contribute to N-Ras depalmitoylation.

**Figure 1.**
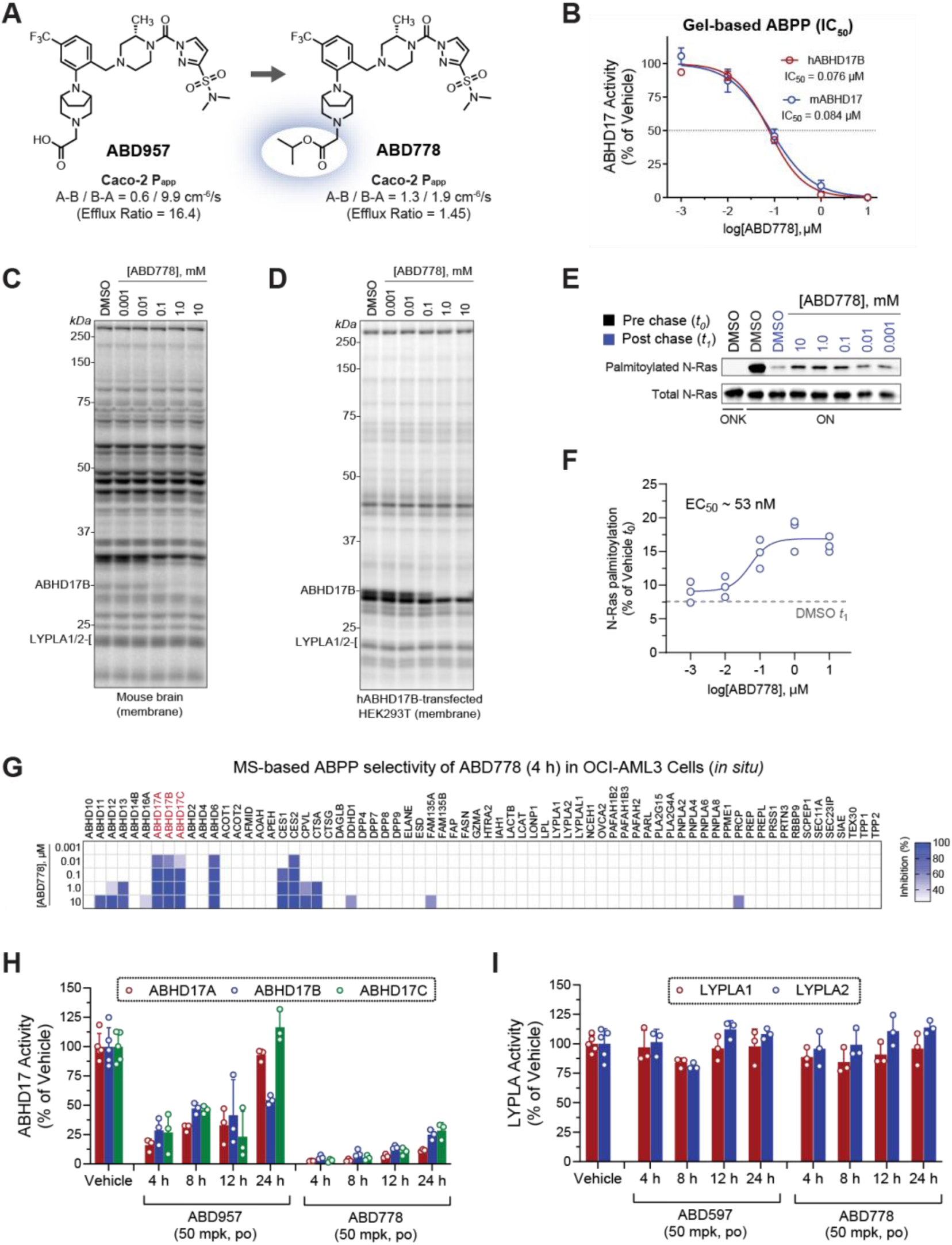
Discovery and characterization of ABD778 as an ABHD17 inhibitor with *in vivo* activity. A, Structures of ABD957 and ABD778 and corresponding Caco-2 permeability data highlighting improved passive permeability and reduced efflux of ABD778. **B**, Half maximal inhibitor concentration (IC_50_) curves for ABD778 against endogenous mouse ABHD17 and recombinantly expressed human ABHD17B was measured by gel-based activity based proteomic profiling (ABPP). Data are presented as mean values ± S.E.M (n = 2-3). **C,D,** Competitive gel-based ABPP profiling of ABD778 (0.001–10 μM) in mouse brain membrane (panel C) and in HEK293T cells transduced with a cDNA encoding human ABHD17B (panel D) showing selective inhibition of ABHD17 over FP-Rh-labeled mouse and human serine hydrolases. **E**, ABD778 partially inhibits N-Ras depalmitoylation. Representative gel measuring N-Ras^G12D^ palmitoylation by 17-ODYA in the presence of varying concentrations of ABD778 (upper panel). N-Ras^G12D^ was immunoprecipitated via GFP and the degree of palmitoylation visualized by rhodamine azide attached via copper(I)-catalyzed click chemistry to the alkyne of 17-ODYA. Total N-Ras content was measured by western blotting of GFP enrichments (bottom panel). As expected, 17-ODYA labeling was not observed in OCI-AML3/ONK cells in which the N-Ras hypervariable domain (HVR) was replaced by the K-Ras4b HVR. **F**, Calculated IC_50_ values for ABD778 stabilization of N-Ras^G12D^ palmitoylation as measured in **e** (n = 3 per group). **G,** *In situ* MS-based ABPP data from OCI-AML3 cells that were exposed to ABD778 (0.001–10 μM) for 4 hours demonstrating ABHD17A/B/C inhibition at low nanomolar concentrations and selectivity across depalmitoylases LYPLA1, LYPLA2 and ABHD10. Data plotted represent mean competition from three biological replicates. **H**,**I**, *In vivo* target engagement for ABHD17A/B/C (panel H) and LYPLA1/2 (panel I) following ABD957 or ABD778 administration. C57Bl/6 mice were dosed with vehicle, ABD957 (50 mg/kg, po) or ABD778 (50 mg/kg, po). Spleen tissue was collected 4, 8, 12 and 24 hours after compound administration and analyzed by targeted MS-ABPP using FP-biotin enrichment of serine hydrolase enzymes and parallel reaction monitoring (PRM) to detect and quantify unique diagnostic peptides from each ABHD17 and LYPLA enzyme. ABD778 provided near complete and sustained blockade of ABHD17 enzymes over 24 hours whereas ABD957 showed transient inhibitory activity. Both compounds maintained selectivity over LYPLA1 and LYPLA2. Data plotted represent the median from biological replicates, and error bars represent S.D. (n = 3-5).

We next evaluated the activity of ABD778 on N-Ras palmitoylation in AML cells. OCI-AML3/ON and OCI-AML3/ONK are isogenic OCI-AML3 sublines that are dependent on either green fluorescent protein(GFP)-labeled mouse N-Ras^G12D^ (ON cells) or a similar fusion protein in which the N-Ras HVR was replaced by the K-Ras4b HVR (ONK cells) for proliferation.^5^ Pulse-chase labeling with an alkynylated (clickable) palmitate analog 17-octadecynoic acid (17-ODYA) revealed selective inhibition of N-Ras^G12D^ depalmitoylation by ABD778 in OCI-AML3/ON cells at a similar EC_50_ value as previously observed for ABD957 (**Figures 1E, 1F**).^5^ We additionally confirmed that ABD778 potently inhibited all ABHD17 isoforms in OCI-AML3 cells while maintaining a similar selectivity profile as observed in cell lysates (**Figure 1G**).

To evaluate the suitability of ABD778 for *in vivo* studies, we dosed C57Bl/6 mice with vehicle, ABD957 (50 mg/kg, og), or ABD778 (50 mg/kg, og) and measured ABHD17 target engagement 4, 8, 12 and 24 hours later in spleen tissue using targeted MS-ABPP. ABD778 produced maximal inhibition of all ABHD17 isoforms at 4 hours and maintained >50% occupancy over the 24 hour time-course (**Figure 1H**). In contrast, ABD957 only partially inhibited ABHD17 enzymes at early time points (4-8 hours) and this effect was not sustained at later time points (24 hours; **Figure 1H**). Consistent with their ABPP selectivity profiles *in vitro*, neither ABD957 or ABD778 inhibited LYPLA1 and LYPLA2 (**Figure 1I**). While ABD778 could be converted through esterolysis to ABD957, we observed limited evidence for this transformation *in vivo* (**Figure S1C**). This finding and the potent inhibition of ABHD17 enzymes by ABD778 *in vitro* suggest that ABD778 itself, rather than ABD957 derived from ABD778 hydrolysis, is likely the active drug in mice. Together, these data identify ABD778 as a selective and *in vivo*-active pan-ABHD17 inhibitor.

### ABD778 inhibits the growth of *NRAS*-mutant leukemia cells and is synergistic with PD901 *in vitro*

ABD778 reduced the growth of OCI-AML3 and of *NRAS* mutant HL-60 cells, but not of the *KRAS*-mutant AML cell lines NB4 and SKM1 (**Figure 2A**). By contrast, PD901 non-selectively suppressed the proliferation of both *NRAS-*and *KRAS*-mutant AML cells (**Figure 2B**). Exposing OCI-AML3/ON and ONK cells to ABD778 confirmed that growth inhibition is dependent on the N-Ras HVR and correlated with reduced ERK phosphorylation (**Figure S2**). We hypothesized that co-treatment with PD901 might augment the genotype-specific activity of ABD778. Indeed, ABHD17 and MEK inhibition synergistically inhibited the growth of *NRAS*-mutant, but not *KRAS*-mutant, AML cells as assessed by the Bliss independence method (**Figure 2C, Figure S3A**).

**Figure 2.**
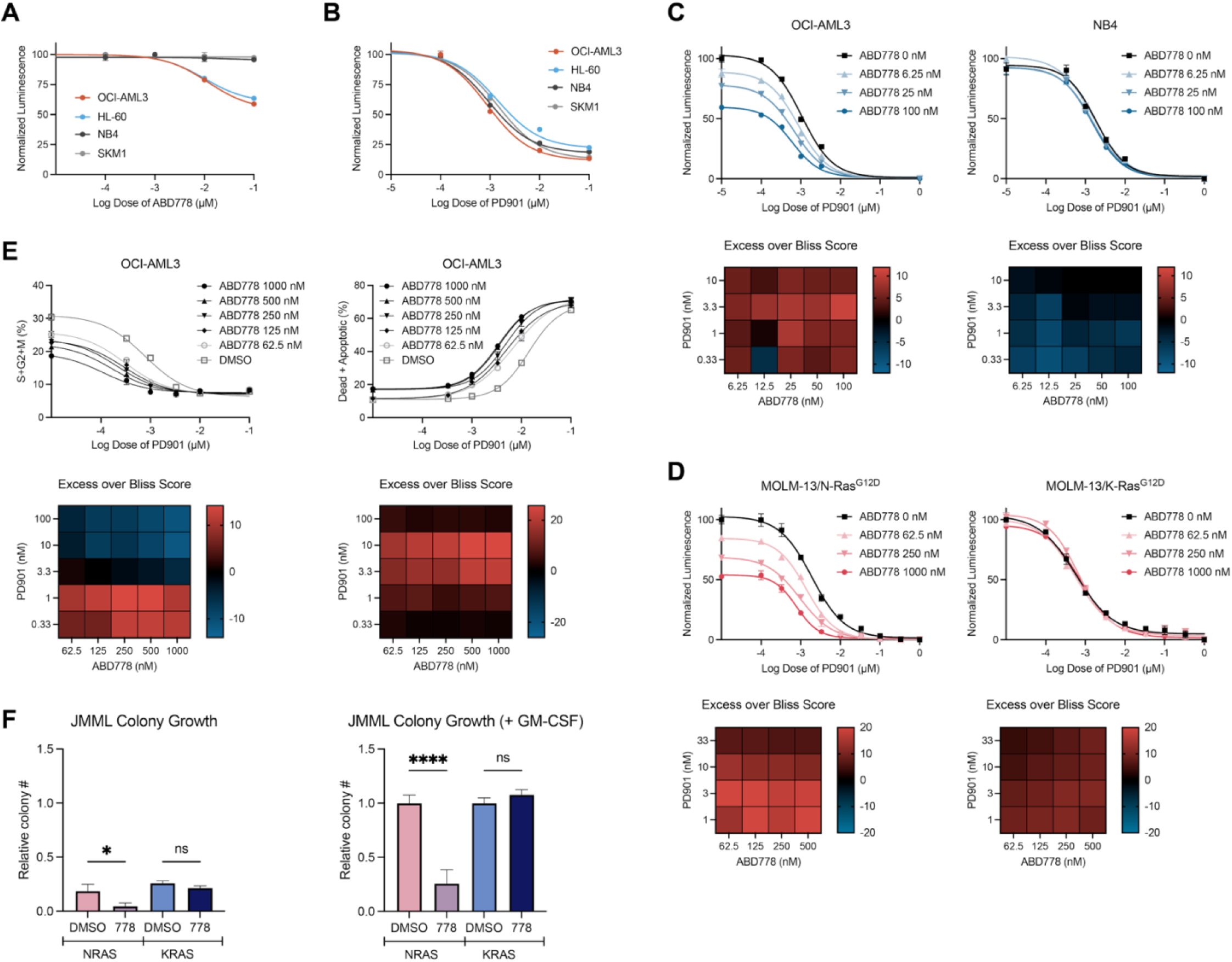
Selective activity of ABD778 and synergy with PD901 in *NRAS*-mutant myeloid leukemias. A, B, Proliferation of *NRAS* mutant OCI-AML3 and HL-60 cells and of *KRAS* mutant NB4 and SKM1 cells were assessed 72 hours after exposure to ABD778 (panel A) or PD901 (panel B) using Cell Titer-Glo. **C**, Bliss independence analysis of proliferation over a broad range of ABD778 and PD901 concentrations in OCI-AML3 (left) and NB4 (right) cells. **D**, Bliss independence analysis of proliferation in MOLM-13/N-Ras^G12D^ (left) and MOLM13/K-Ras^G12D^ (right) cells that were incubated with doxycycline (Dox) for 24h prior to drug treatment. **E**, Bliss independence analysis of cell cycling (left) and apoptosis (right) in OCI-AML3 cells treated with PD901 and ABD778. Heatmaps in panels **C-E** display calculated synergy scores from strongly positive (red) to negative (blue). The data presented in panels **A–E** were generated in triplicate and were replicated in at least two additional independent experiments. **F**, CFU-GM colonies were grown in methylcellulose medium from JMML patient samples harboring *NRAS* or *KRAS* mutations (n = 3 of each genotype) with and without ABD778 (1 µM). Cytokine independent CFU-GM colony growth is shown on the left and the number of CFU-GM colonies observed in cultures containing a saturating dose of GM-CSF (10 ng/mL) is shown on the right. Colony growth was normalized to 100% (1.0) for *NRAS*-mutant and *KRAS*-mutant patient samples grown in GM-CSF, respectively. Note that the GM-CSF markedly augmented CFU-GM growth in JMML cells of both genotypes and that ABD778 robustly suppressed the growth of *NRAS* mutant leukemias.

We previously reported a “switchable” model in which exposing *FLT3*-*ITD* mutant MOLM-13 AML cells to quizartinib abrogates oncogenic FLT3 receptor tyrosine kinase signaling and renders these cells dependent on exogenous doxycycline-regulated Ras oncoprotein expression for proliferation and survival.^31,32^ In this isogenic system, ABD778 reduced the growth of cells engineered to express N-Ras^G12D^ (MOLM-13/N-Ras^G12D^), but not K-Ras^G12D^ (MOLM-13/K-Ras^G12D^) (**Figure S3B**). By contrast, MOLM-13/N-Ras^G12D^ and MOLM-13/K-Ras^G12D^ cells exhibited similar sensitivities to PD901 (calculated 50% inhibitory concentration (IC_50_) values 1.9 nM and 0.6 nM, respectively) (**Figure S3C**). Bliss independence analysis revealed mutant *NRAS* genotype-selective growth inhibition in MOLM-13/N-Ras^G12D^ cells over a range of ABD778 and PD901 concentrations (**Figure 2D**). Additional analyses showed that co-treatment with ABD778 augmented PD901-induced cell cycle arrest and apoptosis in OCI-AML3 and MOLM-13/N-Ras^G12D^ cells in a dose-dependent and Ras isoform-selective manner (**Figure 2E, Figure S3D**).

JMML is an aggressive myeloproliferative neoplasm initiated by mutations in *NRAS, KRAS,* and other genes that constitutively increase Ras signal output.^33,34^ Cytokine-independent colony-forming unit granulocyte-macrophage (CFU-GM) progenitor growth and hypersensitivity to low concentrations of granulocyte-macrophage colony stimulating factor (GM-CSF) in methylcellulose cultures are cellular hallmarks of JMML that are also observed in bone marrow (BM) cells from *Kras^G12D^* and *Nras^G12D^* knock-in mice.^35–37^ As expected, primary *KRAS-*and *NRAS-*mutant CD34-positive JMML cells exhibited cytokine-independent CFU-GM colony growth that was augmented by the addition of a saturating dose of GM-CSF (**Figure 2F, Table S1**). ABD778 significantly reduced both cytokine-independent and GM-CSF-stimulated CFU-GM colony formation from JMML patient samples with *NRAS* mutations but did not alter myeloid progenitor growth from *KRAS*-mutant JMMLs (**Figure 2F**).

### Combined ABHD17 and MEK inhibition durably represses MAPK pathway signaling and blunts feedback reactivation in *NRAS*-mutant cells

Western blot and phospho-flow cytometric analyses showed that ABD778 and PD901 cooperatively reduced phosphorylated ERK (pERK) levels in *NRAS*-mutant OCI-AML3 and HL-60 cells after 4 hours, but not in *KRAS*-mutant NB4 or SKM1 cells (**Figures 3A, 3B, Figure S4**). Phosphorylated S6 (pS6) levels were similarly decreased in OCI-AML3 cells that were exposed to ABD778 and PD901 for 4 hours, but not in HL-60, NB4, or SKM1 cells (**Figures 3A, 3B, Figure S4**).

**Figure 3.**
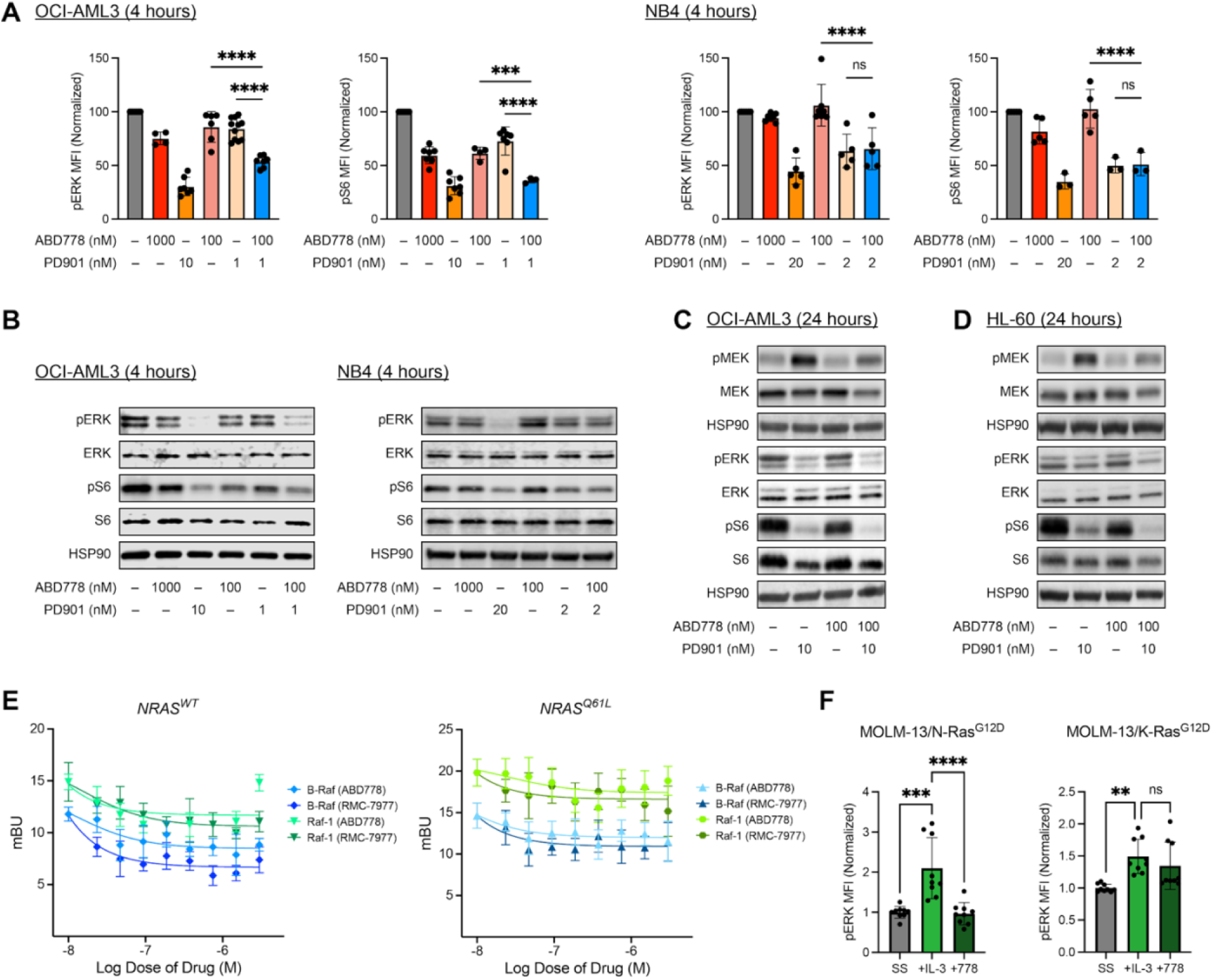
Co-inhibition of MEK and ABHD17 activity cooperatively reduces MAPK pathway activation. **A**, Phosphorylated ERK (pERK) and S6 (pS6) levels were measured by phospho-flow cytometry in OCI-AML3 (left) and NB4 (right) cells that were exposed to ABD778 and/or PD901 for 4 hours at the doses shown. Mean fluorescence index (MFI) values were normalized to 100% of the DMSO control and pooled for statistical analysis. n = 4-11; *** -p < 0.001; **** - p < 0.0001. **B,** pERK and pS6 levels were measured by Western blotting in OCI-AML3 (left) and NB4 (right) cells that were exposed to DMSO, ABD778, PD901, or both drugs for 4 hours. **C**,**D**, pMEK, pS6, and pERK levels were measured by Western blotting in OCI-AML3 (panel **C**) and HL-60 (panel **D**) cells that were exposed to DMSO, ABD778, PD901, or both drugs for 24 hours. **E**, HEK-293T cells were transfected with a fluorescent acceptor HaloTag-NRAS construct (WT left, Q61L right) along with either a Raf-1-NanoLuciferase or B-Raf NanoLuciferase fluorescent donor construct. Corrected ratios of acceptor / donor fluorescent emission (mBU) 1 hour after exposure to ABD778 or the tri-complex RAS(ON) inhibitor RMC-7977 (positive control) are plotted against drug doses. **F**, pERK levels were measured using phospho-flow cytometry in MOLM-13/N-Ras^G12D^ (left) and MOLM-13/K-Ras^G12D^ (right) cells after 24 hours in serum-free media + DMSO (SS), 24 hours in serum-free media + DMSO followed by IL-3 stimulation (+IL-3), and 24 hours in serum-free media + 100nM ABD778 followed by IL-3 stimulation (+778). n = 9; ** - p < 0.01; *** - p < 0.001; **** - p < 0.0001. The data presented were generated in (panels A and F) or are representative of (panel B-E) at least three independent experiments.

*KRAS*-mutant adenocarcinomas restore oncogenic MAPK signaling upon extended exposure to PD901, which is characterized by upregulated Raf kinase activity, elevated pMEK levels, and ERK reactivation.^38–40^ We observed durable repression of ERK and S6 phosphorylation in OCI-AML3 and HL-60 cells that were treated with ABD778 and PD901 for 24 hours in comparison to PD901 alone (**Figures 3C, 3D**). As expected, pMEK levels increased from baseline in OCI-AML3 and HL-60 cells that were exposed to PD901 for 24 hours. ABD778 treatment blunted this increase in pMEK levels (**Figures 3C, 3D**). Consistent with these data, live cell NanoBRET proximity assays showed that ABD778 treatment decreased the interaction between either B-Raf and Raf-1 and WT N-Ras or N-Ras^Q61L^ in transfected HEK-293T cells (**Figure 3E**).

To ask if ABHD17 inhibition interferes with MAPK activation downstream of cytokine growth factor receptors and mutant N-Ras, we exposed serum-starved MOLM-13/N-Ras^G12D^ and MOLM-13/K-Ras^G12D^ cells to ABD778 for 24 hours and then stimulated them with interleukin-3 (IL-3). ABD778 selectively suppressed ERK activation in MOLM-13/N-Ras^G12D^ cells (**Figure 3F**). STAT5 is recruited to the IL-3 receptor and phosphorylated in response to IL-3 stimulation.^41^ As expected, we observed robust STAT5 activation in both MOLM-13/N-Ras^G12D^ and MOLM-13/K-Ras^G12D^ cells after IL-3 stimulation that was unaffected by ABD778 treatment (**Figure S5**).

### *In vivo* efficacy of MEK and ABHD17 inhibition in primary *Nras^G12D^* AMLs

Before initiating preclinical trials in mice, we asked if PD901 alters ABD778 blood levels or ABHD17 target engagement in peripheral tissues. Implementing a more effective protocol for suspending PD901 allowed us to use a lower dose than in previous preclinical trials (2.5 versus 5 mg/kg day).^42–44^ Spleen, kidney and blood samples were collected 2, 6 and 24 hours after administration of vehicle, ABD778 (60 mg/kg, og), PD901 (2.5 mg/kg, og) or both compounds dosed 1 hour apart. ABD778 and PD901 levels in the blood were comparable at each time point measured whether dosed individually or in combination (**Figure S6A, S6B**). Accordingly, ABD778 dosed alone or in combination with PD901 produced robust and similar target engagement across ABHD17 isoforms in spleen and kidney as determined by ABPP (**Figure S6C, S6D**). Extended single-agent ABD778 treatment was well tolerated in WT mice with no evidence of hematologic toxicity **(Figure S7**).

We next used an established PD assay to ask if treatment might suppress cytokine-induced ERK activation *in vivo*.^44,45^ In this experiment, WT mice were treated with control vehicle, ABD778, PD901, or this combination for 4 days and BM cells were isolated 2, 6, and 24 hours after the final drug dose(s). These cells were exposed to a saturating dose of GM-CSF *ex vivo* and pERK levels were measured by flow cytometry in Mac1-positive (Mac1^+^) myeloid cells (**Figure S8A**).^44,45^ Mac1^+^ BM cells isolated 2 and 6 hours after the last dose of either PD901 or the ABD778/PD901 combination had similar reductions in ERK activation in response to GM-CSF stimulation *ex vivo*. However, while BM cells collected from mice 24 hours after the last dose of PD901 showed robust ERK reactivation, this adaptive response was suppressed by combination treatment (**Figure 4A**). As in AML cell lines, ABD778 treatment did not alter STAT5 activation in response to GM-CSF (**Figure S8B**). Based on these data, we hypothesized that co-treatment with ABD778 and PD901 would show greater anti-leukemia activity in preclinical models of *NRAS*-mutant AML than either single agent due to deeper and more durable inhibition of Ras/MAPK signaling.

**Figure 4.**
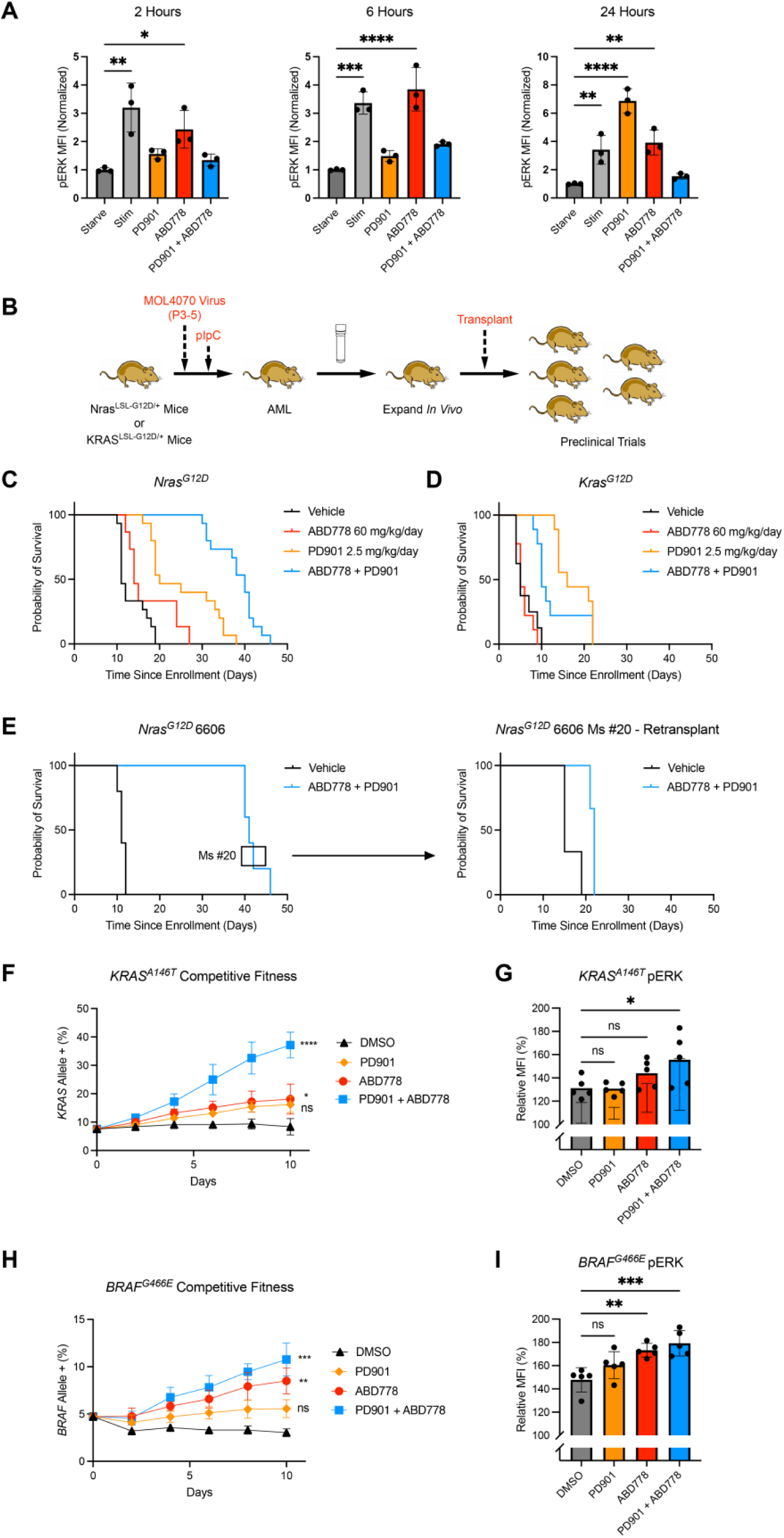
Pharmacodynamics and *in vivo* efficacy of ABD778, PD901, and combination treatment in primary *Nras^G12D^* and *Kras^G12D^* AMLs. A, Phospho-flow analysis of ERK activation in GM-CSF stimulated Mac1^+^ bone marrow (BM) cells collected from mice treated with vehicle, ABD778 (60mg/kg, po), PD901 (2.5mg/kg, po), or both compounds for 4 days. BM cells were collected at 2, 6, or 24 hours after final dose administration. MFI normalized to unstimulated (Starve) condition at each timepoint. * - p < 0.05; ** - p < 0.01; *** - p < 0.001; **** - p < 0.0001. **B,** Transplantable primary AMLs were generated by injecting neonatal wild-type (WT), *Mx1-Cre; Nras^LSL-G12D/+,^* or *Mx1-Cre; Kras^LSL-G12D/+,^*mutant mice with the MOL4070LTR retrovirus followed by a single dose of polyI-polyC at weaning to induce *Nras^G12D^* or *Kras^G12D^*expression from the endogenous loci as described previously^37,46^. Preclinical trials are performed by expanding cryopreserved leukemia cells *in vivo* and transplanting 1-2 x10^6^ marrow cells into 10-20 congenic recipients^42–44^. **C,** Cumulative probability of survival of mice transplanted with *Nras^G12D^* AMLs 6606, 6695, and 6768 that were treated with the control vehicle (black line), ABD778 (red line), PD901 (yellow line), or PD901 + ABD778 (blue line). GraphPad Prism software was used to generate Kaplan-Meier survival curves (primary endpoint) and statistical significance between individual trial arms was calculated using a two-tailed Log-Rank test. These values are presented in the text. **D**, Cumulative probability of survival of mice transplanted with *Kras^G12D^* AMLs 21B and 63A that received the control vehicle (black line), ABD778 (red line), PD901 (yellow line), or PD901 + ABD778 (blue line). Statistical analysis was performed as described in panel b and significant differences reported in the text. **E**, The left panel shows the survival of 5 recipient mice of *Nras^G12D^* AML 6606 that were treated with ABD778 + PD901 in the initial trial. Leukemia cells collected at euthanasia from the recipient shown in the black box were transplanted into secondary recipients and re-treated with either control vehicle or ABD778 + PD901 (n = 3 mice per group). The right panel shows the survival of these mice. **F-I,** Percentages of mCherry-positive (mCherry^+^) OCI-AML3 cells that were transduced with an *mCherry-KRAS^A146T^* (**F**) or an *mCherry*-*BRAF^G466E^* (**H**) expression vector and tracked over 10 days of exposure to either DMSO (black), PD901 (orange), ABD778 (red), or PD901 and ABD778 (blue). Cells were analyzed and replated with fresh drug added every 48 hours. These data were generated from three independent biological replicates. Day 10 time points were pooled for statistical analysis. Relative pERK levels of *mCherry-KRAS^A146T^* (**G**) or *mCherry*- *BRAF^G466E^*(**I**) transduced OCI-AML3 cells were compared to non-transduced cells within the same drug treatment well. Cells in treatment wells were exposed to DMSO, PD901, ABD778, or both drugs for 4 hours. **** p < 0.0001; *** p < 0.001; ** p < 0.01; * p < 0.05.

Transplantable mouse AMLs generated by retroviral insertional mutagenesis in *Kras^G12D^* and *Nras^G12D^* knock-in mice are robust models for performing controlled preclinical trials for assessing drug efficacy and identifying mechanisms of adaptive resistance.^42–44^ In this system, retroviral integrations in proto-oncogenes such as *Ev11, Myb,* and *Sox4* cooperate with subsequent *Nras^G12D^*or *Kras^G12D^*expression in leukemogenesis.^37,44,46^ This reflects the common pathogenic sequence in human AML whereby *RAS* mutations cooperate with antecedent genetic drivers.^47,48^ For preclinical trials, cryopreserved primary leukemia cells are first expanded in one or more “factory” mice and then transplanted into cohorts of immunocompetent congenic recipients that are treated with vehicle or drug(s) (**Figure 4B**). In previous studies, PD901 had modest single-agent activity in *Kras^G12D^*and *Nras^G12D^* leukemias with only one of 10 independent AMLs acquiring adaptive resistance during continuous drug treatment (Table S2).^42,43^

Having confirmed ABD778 single agent tolerability, minimal *in vivo* drug-drug interactions between ABD778 and PD901, efficient targeted engagement with biochemical activity *in vivo,* we sequentially transplanted five groups of 20 mice each with *Nras-*mutant AMLs 6606, 6695, and 6768 or with *Kras*-mutant leukemias 21B and 63A.^37,42,43^ These recipients were treated continuously with control vehicle, ABD778 (60 mg/kg/day), PD901 (2.5 mg/kg/day), or this combination (n = 5 mice per group) by oral gavage. Kaplan-Meier analysis showed that ABD778 modestly improved the survival of mice transplanted with *Nras-*mutant, but not *Kras*-mutant, AMLs (p = 0.0168) (**Figures 4C, 4D**). Consistent with previous preclinical studies^42,43^, PD901 treatment extended survival in recipients of *Nras^G12D^* and *Kras^G12D^* AMLs by approximately two-fold (p < 0.0001 for both *Nras^G12D^* and *Kras^G12D^* leukemias) (**Figures 4C, 4D, Tables S2, S3**). Co-treatment with ABD778 and PD901 further prolonged survival in mice engrafted with *Nras^G12D^* leukemias from ∼20 to ∼40 days (p < 0.0001 versus vehicle; p < 0.0001 versus PD901 alone) (**Figure 4C**). By contrast, there was no significant difference in the survival of recipients of *Kras^G12D^* leukemias that were assigned to the PD901 and PD901/ABD778 treatment arms (**Figure 4D**). A similar pattern of drug responses seen in the pooled analysis of 100 recipient mice was also evident in each independent AML (**Figure S9**). Single-agent and combination treatment with ABD778 was well tolerated with minimal weight loss that was similar to the vehicle and PD901 groups (**Figure S10**). All recipient mice died with progressive AML characterized by blood leukocytosis, anemia, and splenomegaly (**Figure S10**). We re-transplanted *Nras^G12D^*AMLs isolated from at euthanasia from mice that initially responded to ABD778 and PD901 and retreated them *in vivo*. Secondary recipients had greatly reduced survival in comparison to mice enrolled in the initial trial, verifying intrinsic drug resistance in leukemias that emerged after an initial response to combination treatment (**Figure 4E, Figure S11**).

### Emergent on-mechanism mutations cause resistance to ABD778/PD901

To identify candidate resistance mechanisms, we performed whole exome sequencing (WES) of *Nras^G12D^* AMLs isolated at euthanasia from recipient mice treated with either control vehicle or ABD778/PD901. As in relapsed human AMLs^49^, vehicle and drug-treated leukemias shared multiple “founder” mutations (**Table S4**). Interestingly, leukemia cells isolated from independent recipients of AML 6606 showed a *Nras^G12D^* variant allele frequency (VAF) of ∼80-90% after combination treatment versus ∼60% in mice that received the control vehicle. This likely reflects outgrowth of a pre-existing minor clone with uniparental disomy for the mutant *Nras* allele.^37,50^ WES also uncovered new alterations at variant allele frequencies (VAFs) of 3-17% in *Flt3, Kras, Braf,* and other genes that are frequently mutated in patients treated with signal transduction inhibitors (**Table S4**).^51–54^ The observation that each of these mutations were detected in one of five recipient mice assigned to receive ABD778/PD901 suggests that they were acquired during treatment.

To model the outgrowth of a minor drug-resistant clone, we infected OCI-AML3 cells with lentiviral vectors encoding the respective mutant proteins fused to a mCherry reporter gene at a low multiplicity of infection. Transduced cells were then exposed to vehicle, PD901, ABD778, or the PD901/ABD778 combination. *Kras^A146T^* expression resulted in a significant competitive growth advantage upon drug, but not vehicle, exposure that was most pronounced in cells treated with ABD778 and PD901, with *Kras^A146T^* cells expanding to nearly 40% of the total population after 10 days in culture. Outgrowth was characterized by elevated pERK levels in these cells (**Figures 4F, 4G**). Similarly, *Braf^G466E^* expression increased the competitive fitness of transduced OCI-AML3 cells exposed to ABD778 or ABD778/PD901, which was associated with MAPK pathway activation (**Figures 4H, 4I**). We conclude that primary *Nras^G12D^* AMLs that relapse after combination treatment with ABD778 and PD901 acquire mutations that enhance competitive fitness and restore oncogenic MAPK signaling in the presence of both drugs.

### Dual ABHD17 and MEK inhibition enhances MAPK inhibition and promotes apoptosis in *NRAS*-mutant melanoma cells

Tissue-of-origin and cell fate transcriptional programs are key mediators of Ras oncoprotein activity that modulate responses to chemical inhibitors.^52,55^ To determine whether our findings in *NRAS*-mutant AML models would extend to other cancer histologies, we exposed a panel of *NRAS^Q61X^*-mutant melanoma cell lines to a range of ABD778 and PD901 concentrations. Co-treatment synergistically repressed growth in all tested melanoma lines as assessed by the Bliss independence method and had lower pERK levels 4 hours after exposure to ABD778/PD901 versus either single drug (**Figures 5A, 5B, S12A, S12B**). ABD778 also suppressed feedback MEK phosphorylation across all tested *NRAS*-mutant melanomas at the 4 hour time point. (**Figure 5B**, **S12B)**. To determine the depth and durability of this effect, we performed extended time courses in MM485 and SK-MEL-2 cellls. Of note, SK-MEL-2 cells were relatively insensitive to both ABD778 and PD901 but still showed synergistic growth inhibition in response to combination treatment (**Figure 5A**). Western blot analyses of MM485 and SK-MEL-2 cells revealed pMEK levels that were elevated from baseline 4, 24, and 48 hours after exposure to low or high dose PD901 (**Figure 5C**). By contrast, ABD778 modestly reduced pMEK levels and blunted the increase seen after PD901 treatment. Whereas ABD778 did not inhibit ERK activity, co-treatment with PD901 attenuated rebound ERK and RSK phosphorylation observed 24 and 48 hours after PD901 treatment. These biochemical effects of combination treatment were associated with cleaved PARP induction at lower doses of PD901. Altogether, data from *NRAS*-mutant leukemia and melanoma cells support the working model shown in Figure 5D whereby ABD778 enhances the cytostatic and pro-apoptotic activity of PD901 by reducing oncogenic N-Ras signal output and blunting adaptive feedback responses in *NRAS*-mutant cancer cells.

**Figure 5.**
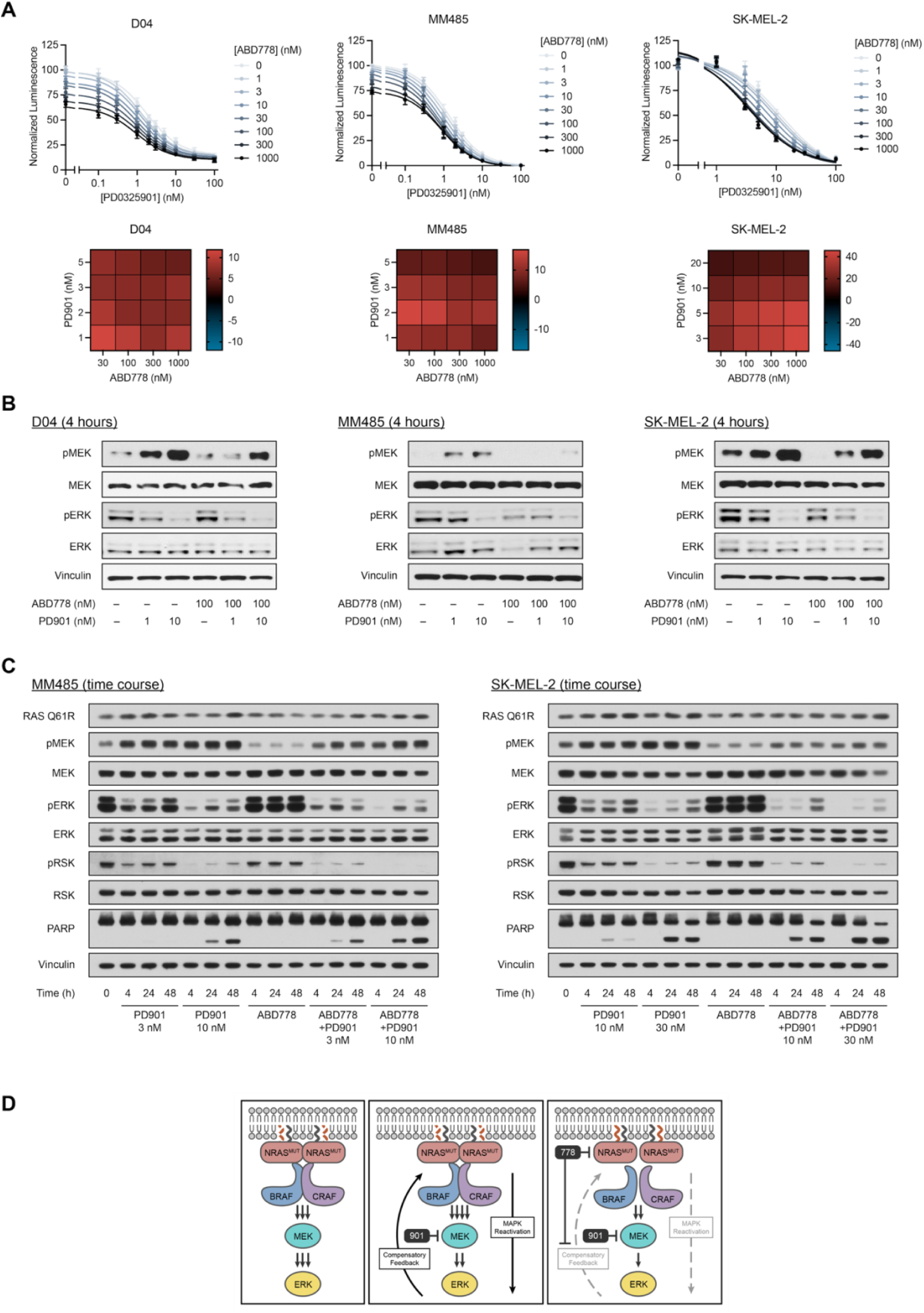
ABD778 enhances PD901-induced growth inhibition and blunts feedback reactivation in *NRAS*-mutant melanoma cell lines. A, Proliferation of *NRAS*-mutant D04, MM485, and SK-MEL-2 melanoma cells was measured after exposure to ABD778 and PD901 using Cell Titer-Glo. Bliss independence analysis is displayed as heatmaps of calculated synergy scores from strongly positive (red) to negative (blue). **B.** pERK and pMEK levels were measured by Western blotting in D04, MM485, and SK-MEL-2 cells that were exposed to DMSO, ABD778, PD901, or both drugs for 4 hours. **C.** pERK, pMEK, pRSK, and PARP cleavage were measured by Western blotting in MM485, and SK-MEL-2 cells that were exposed to DMSO, ABD778, PD901, or both drugs for 4, 24, and 48 hours at the indicated drug doses. Cleaved PARP appears as band below the intact protein. The data presented in **A-C** are inclusive or representative of at three independent experiments. **D,** Working model of synergistic repression of MAPK signaling by ABD778 and PD901. <u>Left</u>. Farnesylated (black curved line) and palmitoylated (red curved line) N-Ras oncoproteins at the plasma membrane (PM) in the active, GTP-bound conformation trigger Raf homo- and heterodimer formation and Raf kinase activation. Raf then constitutively activates MEK and MEK phosphorylates ERK. <u>Middle</u>. PD901 allosterically inhibits MEK kinase activity and thereby decreases ERK activation. Over time, this results in loss of negative feedback, which reactivates oncogenic MAPK signaling. <u>Right</u>. By inhibiting ABHD17 enzymes, ABD778 impairs depalmitoylation and mislocalizes N-Ras within the PM. This, in turn, reduces Raf activation by mutant N-Ras and antagonizes compensatory feedback mechanisms, resulting in deep and durable MAPK inhibition.

### ABD778 augments the activity of Ras pathway inhibitors and overcomes adaptive resistance to FLT3 inhibition

We reasoned that the synergistic activity of ABD778 and PD901 in *NRAS/Nras-*mutant leukemia might extend to additional targeted inhibitors of Ras signaling. In a previous preclinical study, the class 1 PI3 kinase inhibitor pictilisib (also known as GDC-0941) failed to significantly extend the survival of recipient mice transplanted with *Nras^G12^*^D^ AMLs 6606, 6695, and 6768.^42^ Interestingly, however, Bliss independence analysis showed synergistic growth inhibition by ABD778 and pictilisib in *NRAS*-mutant OCI-AML3 cells that was not observed in *KRAS*-mutant NB4 cells (**Figure S13A**). Western blot analysis revealed equivalent reduction of Akt phosphorylation in OCI-AML3 and NB4 cells after short-term exposure to pictilisib or ABD778/pictilisib, while ABD778 decreased pERK and pS6 levels in OCI-AML3 cells after combination treatment (**Figure S13B**). These data suggest that ABD778 inhibits the compensatory increase in MAPK signaling reported by others in pictilisib-treated cell lines.^56^

Covalent allele-selective and Ras tri-complex inhibitors are promising new therapies for patients with *RAS*-mutant cancers.^26,57–59^ Sotorasib is a FDA-approved K-Ras^G12C^ inhibitor that is also active in cancer cell lines harboring *NRAS^G12C^* mutations.^25^ As expected,^32,60^ sotorasib inhibited the growth of isogenic MOLM-13 cells expressing either N-Ras^G12C^ or K-Ras^G12C^. ABD778 and sotorasib had synergistic anti-proliferative activity that was selective for MOLM-13/N-Ras^G12C^ cells and was associated with reduced pERK levels (**Figure S14**). RMC-6236 (daraxonrasib)^26^ is a pan-Ras tri-complex inhibitor that similarly showed genotype-selective synergistic activity with ABD778 and cooperatively repressed pERK levels in MOLM-13/N-Ras^G12D^ cells (**Figures 6A, 6B**).

**Figure 6.**
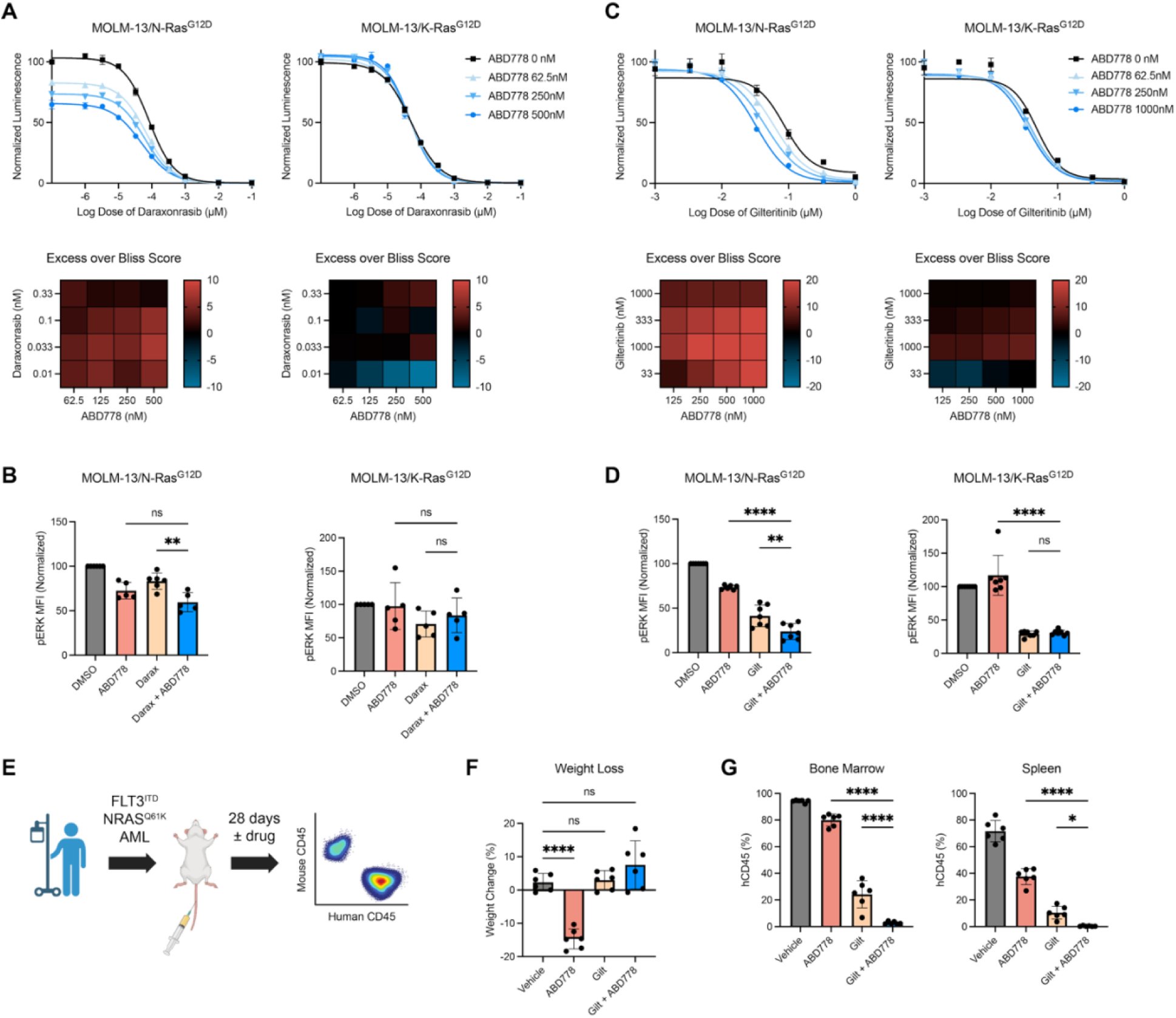
ABD778 cooperates with Ras tricomplex and FLT3 inhibitors to suppress the growth of *NRAS*-mutant AML cells. A, Bliss independence analysis of MOLM-13/N-Ras^G12D^ (left) and MOLM13/K- Ras^G12D^ (right) cells that were treated with doxycycline (Dox) for 24 hours to induce N-Ras^G12D^ or K-Ras^G12D^ expression were exposed to a range of ABD778 and RMC-6236 doses. **B,** pERK levels were measured by phospho-flow cytometry in MOLM13/N-Ras^G12D^ (left) and MOLM13/K-Ras^G12D^ (right) cells that were exposed to the ABD778 and/or RMC-6236 doses shown for 4 hours (n = 6-14). **C,** Bliss independence analysis of MOLM- 13/N-Ras^G12D^ and MOLM13/K-Ras^G12D^ cells that were treated with a range of ABD778 and gilteritinib doses was performed as described above. **D,** pERK levels were measured by phospho-flow cytometry in MOLM13/N- Ras^G12D^ (left) and MOLM13/K-Ras^G12D^ (right) cells that were exposed to the ABD778 and/or gilteritinib doses shown for 4 hours (n = 6-7). **E,** Schema of AML PDX trial approach. Briefly, NSG mice were transplanted with 1 million human AML PDX cells. After confirming the presence of human CD34+ cells on day 4 post-transplant, recipient mice were treated daily with vehicle, ABD778, gilteritinib, or ABD778 + gilteritinib combination (n = 6 mice per group) for a planned 28 days. Created with https://BioRender.com. **F,** Mice in all four treatment cohorts were euthanized at days 21-22 due to weight loss in mice assigned to ABD778 treatment. **** p < 0.0001. **G,** The percentage of human hematopoietic cells in the spleens and bone marrows of NSG mice were determined at the end of the trial by flow cytometry using an antibody to human CD45. **** p < 0.0001, ** p < 0.001, * p < 0.05.

Gilteritinib is an FDA-approved treatment for patients with relapsed/refractory *FLT3*-mutant AML and is now used in many front-line treatment protocols.^61,62^ In patients who enter remission, relapse after gilteritinib treatment is frequently characterized by out-growth of resistant clones harboring both the *FLT3* mutation identified at diagnosis and a secondary oncogenic *NRAS* mutation.^9^ To model this clinical situation *in vitro*, we exposed MOLM-13/N-Ras^G12D^ and MOLM-13/K-Ras^G12D^ cells to a range of gilteritinib concentrations with or without doxycycline. As expected^9^, N-Ras^G12D^ expression induced gilteritinib resistance (**Figure S15**). Co-treatment with ABD778 selectively resensitized MOLM-13/N-Ras^G12D^ cells to gilteritinib as assessed by Bliss Independence analysis, which correlated with reduced pERK levels (**Figures 6C, 6D**).

We next injected an aggressive PDX isolated from an AML patient with a *FLT3*-*ITD* mutation who relapsed after gilteritinib treatment with a secondary *NRAS^Q61K^* mutation into NSG mice and treated them with vehicle, ABD778, gilteritinib, or both drugs (**Figure 6E**). This trial was terminated after 3 weeks due to morbidity and weight loss in mice assigned to ABD778 treatment (**Figure 6F**). At euthanasia, the spleens and BM of mice in the vehicle group were extensively infiltrated with human CD45^+^ AML cells (**Figure 6G**). ABD778 treatment modestly reduced leukemia growth in both organs, with gilteritinib showing more potent activity. Combination treatment profoundly reduced the percentage of PDX cells in the BM of mice that received both drugs and eliminated detectable human cells from the spleen (**Figure 6G**).

## DISCUSSION

Oncogenic Ras proteins were historically viewed as “undruggable” due to their structural features, low nanomolar affinity for guanine nucleotides, and high intracellular GTP levels.^63^ The landmark discovery and subsequent clinical development of covalent K-Ras^G12C^ inhibitors radically changed this view, and multiple isoform-selective and pan-Ras inhibitors are either FDA-approved or undergoing clinical testing.^25,58,64^ Similarly, the disappointing efficacy of farnesyl transferase inhibitors led many in academia and biopharma to abandon Ras post-translational processing as a viable target for drug discovery. Our data demonstrating synergistic activity of dual ABHD17 and MEK inhibition in AML and melanoma models harboring oncogenic *NRAS/Nras* mutations thus opens a potential new avenue for treating *NRAS* mutant cancers.

Mechanistically, we show that ABD778 potently inhibits ABHD17 enzymes, reduces N-Ras depalmitoylation, and partially suppresses constitutively elevated pERK levels in *NRAS*-mutant cancer cells. Accordingly, ABD778 reduced the *in vitro* growth of AML and melanoma cell lines and JMML patient samples in a genotype-selective manner and exhibited potent synergy with PD901. Biochemical and BRET analyses showed that ABD778 reduces oncogenic N-Ras signal output and blunts the adaptive feedback response of *RAS*-mutant cancer cells to PD901. N-Ras^G12D^ is mis-localized within the plasma membrane in response to ABD957, a tool compound that is structurally similar to ABD778 (**Figure 1A**).^5^ We speculate that this, in turn, reduces N-Ras-GTP binding to Raf and the subsequent formation of active Raf-MEK dimers, which augments MAPK pathway inhibition by PD901. ABD778 also selectively and synergistically inhibited the growth of *NRAS*-mutant AML cell lines in combination with mutant allele-specific and pan-Ras tri-complex inhibitors. These data raise the intriguing question of whether ABHD17 inhibition suppresses adaptive MAPK pathway reactivation after exposure to these Ras inhibitors.

Co-treatment with ABD778 and PD901 significantly extended the survival of recipient mice transplanted with congenic *Nras^G12D^,* but not *Kras^G12D^,* leukemias. We detected similar by-pass mutations in murine *Nras^G12D^* AMLs after ABD778/PD901 treatment as those seen in patients with lung adenocarcinomas that progressed after an initial response to K-Ras^G12C^ inhibitor treatment.^65^ Thes data point to the rewiring of Ras pathway activity as a more common resistance mechanism than mutational disruption of drug binding to primary targets (e.g., ABHD17 enzymes, MEK). The *Kras^A146T^* mutation is notable because A146T is less activating than G12D in myeloid lineage cells and other tissues,^52^ yet emerged after treatment with ABD778 and PD901. Taken together, the potent *in vivo* activity of combination treatment, our biochemical data, and the spectrum of emergent resistance mutations further support a mechanism whereby ABD778 augments MAPK pathway inhibition and thereby shapes how *NRAS*-mutant cancers develop adaptive resistance to restore oncogenic signaling.

We believe that these studies validate N-Ras post-translational processing as a therapeutic target in cancer and should accordingly stimulate future mechanistic and translational investigation of the N-Ras palmitoylation cycle. For example, our findings indicate that the sub-chronic inhibition of ABHD17 enzymes is well-tolerated in mice, suggesting that potential effects on additional palmitoylation substrates of these enzymes^5^ do not lead to severe health impairments. Nonetheless, how ABHD17 inhibitors impact the palmitoylation and function of other dynamically palmitoylated proteins^66–71^ warrants further investigation. Inhibiting PAT enzymes or interacting proteins such as the RAB27B GTPase are possible alternative therapeutic strategies for treating *NRAS*-mutant cancers that are supported by genetic, biochemical, and phenotypic data.^3,6,21,72^

AML, melanoma, and other aggressive cancers that initially respond to signal transduction inhibitors or other single targeted agents almost invariably develop adaptive resistance that drives cancer regrowth.^9,65,73^ Accordingly, there is a compelling rationale for implementing effective combination regimens using drugs with complementary mechanisms of action. Promising efficacy data from clinical trials of Ras and epidermal growth factor receptor inhibition in colorectal cancer support implementing therapeutic strategies that anticipate and target adaptive resistance mechanisms.^73^ Our studies of ABHD17 and MEK inhibition illustrate how this principle might be exploited to enhance therapeutic index in AML, melanoma, and other *NRAS*-mutant malignancies by co-targeting Ras post-translational processing and constitutive MAPK pathway activation.

## Acknowledgments

Research reported in this publication was supported by NIH grants R01CA193994, R01CA231991, R35CA231991, U54CA196519, R37CA266550, K08CA256489, and T32CA128583, and by the West Charitable Trust and St. Baldrick’s Foundation Cody Thompson Memorial Hero Fund. MD is a Damon Runyon-St. Jude Pediatric Cancer Research Fellow supported by Damon Runyon Cancer Foundation and St. Jude Children’s Research Hospital fellowship award (SJP-09-25). KS is an American Cancer Society Research Professor and CCS is a Damon Runyon-Richard Lumsden Foundation Clinical Investigator supported (in part) by the Damon Runyon Cancer Research Foundation (CI-99-18) and is a Leukemia and Lymphoma Society Scholar in Clinical Research. We are grateful to our colleagues Jonathan Ostrem, and Kevan Shokat for helpful discussions.

## Author Contributions

M.D., B.J.H., T.W., M.J.N., K.S. and B.F.C. conceived the project and wrote the paper. M.J.N. led efforts to develop and characterize ABD778 with assistance from N.N., D.M.H., R.M.S, C.G., K.S., T.A., E.R., B.X. and J.R.R. and M.M.D. T.W. and R.M.S. performed N-Ras palmitoylation assays. M.D., B.H., C.B., M.T., J.Y., A.C.B., K.A.T., M.A., C.J.L., M.H., and K.T. performed proliferation, N-Ras signaling, competitive growth assays, Bliss Independence analysis, molecular cloning, and cell transduction. C.B., M.H, and K.T. executed the preclinical trials in *Nras^G12D^* and *Kras^G12D^*leukemias and analyzed tissues from recipient mice. H.H. and E.S. performed the colony growth assays of JMML patient samples. J.V. and M.A. carried out BRET experiments. M.S., R.F., E.T.G., and F.M. performed proliferation and N-Ras signaling assays in melanoma lines. C.C.S. generated the PDX model that S.A., J.R., and E.S. used to test the efficacy of ABD778 and PD901. Y.L. and his lab performed exome sequencing and Y.L. collaborated with B.J.H. and M.D. to analyze these data. M.D., B.J.H., T.W., M.J.N., K.S. and B.F.C. performed data analysis and visualization.

## Competing interests

Nhi Ngo, Dylan M. Herbst, Radu M. Suciu, Carlos Guijas, Kimia Sedighi, Taylor Andalis, Elysia Roche and Boer Xie are current employees of Lundbeck. Micah J. Niphakis was employed by Lundbeck when this work was performed and is no longer employed there. The authors declare no other competing interests.

## Methods

### ABD778 synthesis

The Extended Data sections includes a description of the synthesis and characterization of ABD778 with the structures of intermediate compounds.

### Caco-2 permeability assay

Compound permeability was determined using a Caco-2 cell monolayer bidirectional transport assay coupled to LC-MS/MS analysis. Caco-2 cells (50 µL at 6.86×10⁵ cells/mL) were dispensed into each well of a 96-well HTS Transwell plate and cultured for 14-18 days.

Test compound solutions were prepared as 1.0 mM stock solutions in DMSO and diluted in HBSS buffer (10 mM HEPES, 2% BSA, pH 7.4) to 5.0 µM. To measure apical-to-basolateral (A→B) transport, 75 µL of test compound solutions were added to the Transwell insert (apical compartment) and 235 µL HBSS buffer (10 mM HEPES, 2% BSA, pH 7.4) to the receiver plate (basolateral compartment). For basolateral-to-apical (B→A) transport, 235 µL of test compound solutions were added to the receiver plate (basolateral compartment) and 75 µL of HBSS buffer was added to the Transwell insert (apical compartment). Plates were incubated at 37°C for 2 h. Samples (50 µL) were collected from apical and basolateral compartments and quenched with 4 volumes of cold methanol. Samples were vortexed and centrifuged at 3,220g for 40 min, and supernatants (100 uL) were diluted 1:1 with water for LC-MS/MS analysis.

Sample extracts (3 µL) were injected onto a Waters XSelect HSS T3 C18 column (2.1 x 50 mm, 2.5 µm particle size) coupled to a Waters XSelect HSS T3 C18 guard column (2.1 x 30 mm, 2.5 µm particle size). Mobile phase A consisted of 0.1% formic acid in water, and mobile phase B consisted of 0.1% formic acid in acetonitrile. The gradient started at 5% B for 0.8 min, increased linearly to 100% B over 0.3 min, maintained at 100% B for 0.1 min, before returning to initial conditions. Flow rate was maintained at 650 µL/min.

Analytes were quantified using an API 5500 mass spectrometer equipped with an ESI interface operated in MRM mode. Source parameters were maintained as follows: collision gas = 6 L/min, curtain gas = 30 L/min, nebulizer gas = 50 L/min, auxiliary gas = 50 L/min, temperature = 500°C, and ion spray voltage = +5500V. The quantitative transition for ABD778 was m/z 669.94➔369.21 (DP = 120V, EP = 8V, CE = 40V, CXP = 15V). The quantitative transition for ABD957 was m/z 628.50➔327.30 (DP = 80V, EP = 8V, CE = 32V, CXP = 15V).

Apparent permeability coefficients (Papp) were calculated using the equation:

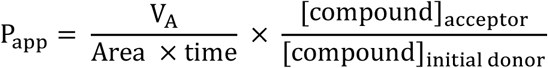

*V_A_* is the acceptor volume, Area is the membrane surface area (0.143 cm²), and time is the transport duration in seconds. The efflux ratio was determined using the following equation:

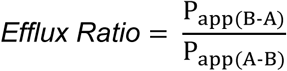

### Cell lines and tissue culture

OCI-AML3 (DSMZ, ACC-582), NB-4 (DSMZ, ACC-207), HL60 (DSMZ, ACC-3), SKM1 (DSMZ, ACC-547) cells, and lab-generated MOLM13 lines were grown in RPMI supplemented with 10% fetal bovine serum (FBS), L-glutamine (2 mM), penicillin (100 U ml^−1^), streptomycin (100 μg ml^−1^) and grown at densities between 0.3 x 10^6^ and 2 x 10^6^ cells mL^−1^. HEK293T (ATCC, CRL-3216) were grown in DMEM supplemented with 10% FBS, l-glutamine (2 mM), penicillin (100 U ml−1) and streptomycin (100 μg ml−1). All cells were maintained at 37 °C with 5% CO2.

### Proteome preparation

Tissue proteomes (mouse brain, kidney, and spleen) were prepared by adding each tissue, a 5 mm stainless steel bead, and cold PBS (1.0 mL) to Safe-Lock tubes and homogenizing using the Tissue Lyser II (Qiagen) (30 Hz/sec for 30-60 sec). For spleen and kidney samples, protease inhibitors (Pierce^TM^ Protease Inhibitor Mini Tablets, EDTA-free) were added to the PBS prior to homogenization. Homogenized tissues were then clarified by centrifuging (2500g) at 4 °C and discarding the pellet. Cell proteomes were prepared by sonicating cell pellets in cold PBS. To isolate the membrane fraction of tissues or cells, cell lysates or brain homogenates were subjected to ultra-centrifugation (100,000g for 45 min at 4 °C). Supernatants were removed and membrane pellets were resuspended in PBS using probe sonication. Protein concentrations were determined using Bio-Rad DC protein assay and diluted to desired protein concentration (see below) for subsequent ABPP analysis.

### Gel-based ABPP

Inhibitor potency (IC_50_ values and target engagement) against ABHD17 and other serine hydrolases was determined by competitive gel-based ABPP in mouse brain and hABHD17B-transfected HEK293T membrane proteomes using FP-Rh competition^1^. hABHD17B in a pCMD-SPORT6 vector (MHS1010-202726047, 4748883, Dharmacon) was recombinantly expressed in HEK293T cells.

*In vitro* potency for mABHD17 and hABHD17B was determined by treating mouse brain or hABHD17B-transfected HEK293T membrane proteomes (50 µg, 1.0 mg/mL) with ABD778 (0.001 – 10 µM) or DMSO in triplicate for 30 min at 37 °C and subsequently treating with FP-Rh (1.0 µM) for an additional 30 min at room temperature. After incubation with FP-Rh, reactions were quenched with 4X SDS-PAGE loading buffer and FP-Rh-labeled enzymes were resolved by SDS-PAGE (10% acrylamide). In-gel fluorescence was visualized using a Bio-Rad ChemiDoc^TM^ XRS imager. Fluorescence is shown in gray scale. Quantification of enzyme activities was performed by densitometric analysis using ImageJ software (NIH). Integrated peak intensities were generated for the band corresponding to mABHD17 in mouse brain and hABHD17B in HEK293T proteomes. Note that the identity of the ABHD17 band detected in mouse brain ABPP gels may correspond to any one or more of the ABHD17 isoforms, therefore we have not specified which isoform is being measured. ABHD17 IC_50_ values were calculated through curve fitting semi-log-transformed data (*x*-axis) by non-linear regression with a four-parameter, sigmoidal dose response function (variable slope) in Prism software (GraphPad).

### In situ selectivity profiling by MS-ABPP

For *in situ* treatments, OCI-AML3 cells (5 × 10^6^ cells) were added to 6-well plates at a density of 1.25 × 10^6^ cells ml^−1^. Cells were treated with DMSO or ABD778 (0.001 – 10 µM) in triplicate and incubated for 4 h. Cells were pelleted at 500g by centrifugation, washed with PBS, pelleted again and snap frozen prior to cell lysis, proteome preparation and ABPP.

### MS-based ABPP sample preparation

For *in vitro* MS-ABPP, cell or tissue proteomes (1.0 mg/mL in 0.2 mL of PBS) were treated with inhibitor (0.001-10 mM) or DMSO for 30 min at 37 °C and subsequently labeled with FP-biotin (10 μM) for 1 h at room temperature. Proteomes from *in situ*-treated cell pellets (1.0 mg/mL in 0.2 mL of PBS) or tissues collected from mice following compound treatments (5.0 mg/mL in 0.2 mL of PBS) were directly labeled with FP-biotin as described above.

Following FP-biotin treatment, proteomes were denatured and precipitated using 9:1 acetone/MeOH, resuspended in 0.2 mL of 8 M urea in PBS and 1% SDS, reduced using DL-dithiothreitol (DTT, 10 mM) for 20 min at 55 °C, and then alkylated using iodoacetamide (50 mM) for 30 min at room temperature in the dark. The biotinylated proteins were enriched with PBS-washed streptavidin-agarose beads (50 μL; Thermo Scientific) by rotating at room temperature for 1.5 h in PBS with 0.2% SDS (1.3 mL). The beads were then washed sequentially with 0.5 mL 0.2% SDS in PBS (10×), 1.1 mL PBS (10×) and 1.1 mL DI H_2_O (10×). On-bead digestion was performed using sequencing-grade trypsin (2 μg; Promega) in 2 M urea in PBS for 12–14 h at room temperature (100 μL). The beads were removed using filtration and washed with DPBS (100 µL). Protein digests were acidified by addition of formic acid (10 µL of 100% formic acid) and desalted using SOLAµ^TM^ SPE Plates (HRP 2 mg / 1 mL). Samples were dried by centrifugal evaporation and stored at -80 °C.

For *in vitro* and *in situ* ABPP samples, peptides were labeled with TMTpro reagents using protocols adapted from previous work^2^. Dried samples were reconstituted in 20 µL EPPS buffer, pH 8.5 followed by addition of 5 µL 40 mM TMTpro reagents in acetonitrile for 1 hr. Reactions were quenched with 5 µL 5% hydroxylamine, mixed, desalted, dried by centrifugal evaporation, and stored at -80 °C until analysis.

### MS data acquisition and analysis

#### Parallel reaction monitoring (PRM) for in vivo target engagement

Dry peptide samples were reconstituted in water containing 0.1% formic acid (20 µL) and 10 µL were injected onto an EASY-Spray column (15 cm x 75 µm ID, PepMap C18, 3 µm particles, 100 Å pore size, Thermo Fisher Scientific) using a Vanquish Neo UHPLC (Thermo Fisher Scientific). Peptides were separated over a 15 min gradient of 0 to 40% acetonitrile (0.1% formic acid) and analyzed on an Orbitrap Fusion Lumos (Thermo Fisher Scientific) operated using a parallel reaction monitoring (PRM) method targeting select peptides from ABHD17A, ABHD17B, ABHD17C, LYPLA1 and LYPLA2 as well as additional control peptides to assess sample integrity: PCCA and PC. Selected ions were isolated and fragmented by high energy collision dissociation (HCD) at 30% CE and fragments were detected in the Orbitrap at 15,000 resolution. Further details for the targeted peptides can be found in Table 1.

**Table 1.**
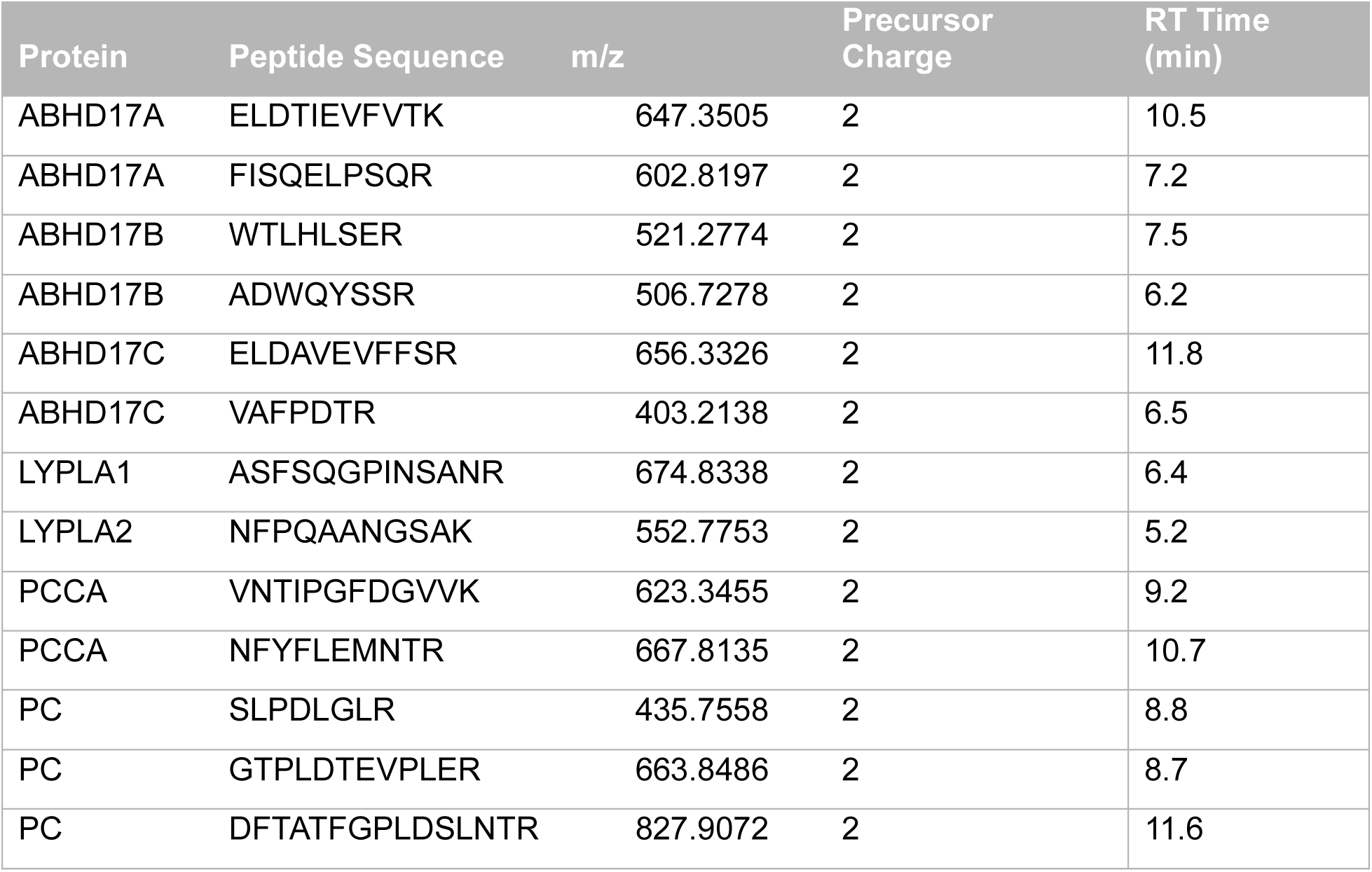

#### MS Acquisition for TMTpro

##### Labeled Peptides

MS was performed using a Thermo Scientific Vanquish Neo and Orbitrap Eclipse system. Peptides were eluted over a 240 min nLC 5-25-45% acetonitrile gradient. For all samples, data were collected in data-dependent acquisition mode over a range from 375–1,500 m/z. Each full scan was followed by fragmentation events for 1.5 sec including real-time search of mouse or human peptides with TMTpro modifications and carbamidomethylation of cysteine and oxidation of methionine for subsequent synchronous precursor selection. Dynamic exclusion was enabled (repeat count of 1, exclusion duration of 60 s) for all experiments.

##### TMTpro Data Analysis

Data analysis was performed using a custom in-house pipeline. Raw files were converted to the indexed mzML format using ThermoRawFileParser v1.4.2^3^ and then searched with Comet v2019.01.5^4^ against a human FASTA database obtained from UniProt ^2^ (reference proteome UP000005640_9606, downloaded on 2020-03-21), and concatenated with a list of non-human contaminant proteins obtained from MaxQuant^5^, and reverse decoys. For the Comet search, TMTPro modifications on lysine and the N-terminus were specified as static modifications, in addition to a static modification for carbamidomethylation at cysteine [+57.021463], and a differential modification for oxidation at methionine [+15.994914]. Further processing was performed using the OpenMS platform v3.0.0^6^ (commit short hash 26292ad). Files were first converted from .pep.xml to .idXML using IDFileConverter, target/decoy information was added using PeptideIndexer, features were extracted with PSMFeatureExtractor, and Percolator v3.05^7^ was run through PercolatorAdapter. Peptide spectrum matches (PSMs) were then filtered with a q-value threshold of 0.01. Further processing was then carried out using custom Python code. For each filtered PSM, MS3 scans were extracted from the original mzML file, and reporter ion intensities were extracted from the most intense peak within a 0.002 Th window of each reporter ion expected mass. Intensity values were then normalized per channel to the mean per-channel summed intensity. PSMs were excluded from downstream calculations if their control sum intensity was less than 5000 x N where N is the number of channels corresponding to the control condition. Intensity values for each PSM were then summed per-channel for each protein, and percent activity values were calculated by dividing each value by the mean intensity across the summed control channels. The mean was then taken across channels associated with the same condition and was reported as a percent activity value for each protein.

###### *In vivo* target engagement and bioanalysis of ABD778 and ABD957

Female C57Bl/6J mice (Jackson) aged 10 weeks at the time of dosing were administered ABD778, ABD957 or PD-901 by oral gavage (10 ml/kg volume). Each compound was prepared fresh on the day of dosing in PEG400:ethanol:PBS (7:2:1) for ABD778 and ABD957 or 0.6% methylcellulose and 0.2% Tween80 in water for PD901. Maximal dispersal of the compound was achieved by bath and probe sonication until a uniform white suspension was formed. For combination studies, PD901 was administered 1 h prior to ABD778. Animals were anesthetized with isoflurane at the indicated time points after compound administration. Blood was collected by cardiac puncture into EDTA microtubes, quenched with 4 volumes of cold acetonitrile and centrifuged at 17,000 x g for 2 minutes prior to collecting the supernatants for analysis. Spleen and kidneys were then removed and rinsed in PBS before freezing in liquid nitrogen. Tissues were stored at -80 C until analysis by ABPP (see below). For pharmacokinetic analysis of ABD778 and ABD957-derived from hydrolysis of ABD778, blood (∼30 µL) was serially collected from the dorsal metatarsal vein and quenched with cold acetonitrile (120 µL) and stored at -80 C until analysis.

###### LC/MS analysis of compound concentrations from whole blood

Compound concentration in acetonitrile-quenched whole blood was determined by ultra-performance liquid chromatography coupled to mass spectrometry (UPLC/MS). The calibration curve samples were prepared using whole blood from naïve mice quenched with 4 volumes of acetonitrile following the same protocol as above and spiked with serial dilutions of the test article, ranging 0.1-5000 ng/mL. Samples were incubated on ice for 60 minutes and then centrifuged at 2,400 *g* at 4 °C for 15 minutes and supernatants were transferred to another tube. 20 mL of supernatant were mixed with 80 mL of water and transferred to a LC vial containing a borosilicated glass insert for UPLC/MS analysis.

Whole blood extracts (10 mL) were injected onto an Agilent 1290 UPLC system equipped with a G7120A pump, a G7167B multisampler and a G1170A column manager (Agilent Technologies, Santa Clara, CA). Chromatographic separation was achieved using an Acquity UPLC BEH C18 Column (2.1 x 50 mm, 1.7 mm particle size, 130 Ǻ) coupled to an Acquity UPLC BEH C18 Column Guard (2.1 x 5 mm, 1.7 mm particle size, 130 Ǻ) (Waters Corporation, Milford, MA). Mobile phase A was composed of water/acetonitrile 95:5 (v/v) and mobile phase B was composed of acetonitrile/water 95:5 (v/v) with 0.1% formic acid added to both mobile phases. The gradient started with 0% B for 0.5 min before being increased linearly to 100% B over 4 min. Afterwards, solvent B was kept at 100% for 1 min, before switching to the initial conditions in 0.1 min. The system was allowed to equilibrate for 1.4 minutes before the next sample injection. Flow rate was kept at 0.6 mL/min and the column temperature was 50 °C.

Analytes were quantified using a 6470 triple quadrupole mass spectrometer equipped with an electrospray Jet Stream source (Agilent Technologies) operated in dynamic multiple reaction monitoring (dMRM) mode. The quantitative and qualitative transitions for each compound were optimized using the authentic standards in the Optimizer software (Agilent Technologies): ABD778 (Quant: 670.3➔369.1, CE=22 eV; Qual: 670.3➔ 240.0, CE= 45 eV); PD901 (Quant: 483.0➔249.0, CE= 22 eV; Qual: 483.0➔ 375.9, CE= 10 eV). The following parameters were kept constant for all transitions: Fragmentor=180, Cell Accelerator Voltage=4, Polarity=Positive. Total cycle time was 500 ms. Source parameters were kept as follows: Dry Gas Temperature=350 °C, Dry Gas Flow=11 L/min, Sheath Gas Temperature=350 °C, Sheath Gas Flow=11 L/min, Nebulizer= 50 psi, Noozle voltage= 1500 V (positive) and Capillary= 3500 V (positive). Compounds concentrations were calculated by interpolating the integrated area under the curve with the calibration curves for each tissue prepared in the same matrix than the samples using the Masshunter Quantitative Analysis Software (Agilent Technologies).

###### Isogenic OCI-AML3/ON and ONK cell lines

A pCDH-LMN–GFP lentiviral vector was obtained by cloning the miR30-PGK-NeoR-IRES–GFP cassette from LMN–GFP^8^ into a pCDH Expression Lentivector (System Biosciences). A miR30-based shRNA targeting human *NRAS* (sense, 5′-CAGGGTGTTGAAGATGCTTTT-3′) was cloned into the vector. The coding sequence of *Nras^G12D^* was cloned downstream of GFP to create a N-terminal GFP-fused N-Ras^G12D^ expression construct with N-Ras endogenous HVR (N-HVR). Alternatively, a chimeric version was cloned where the sequence corresponding to amino acids 166–188 from N-Ras was replaced with that of K-Ras4b (K-HVR). Lentiviral vector production and transduction of OCI-AML3 cells were performed as previously described^1^.

###### Dynamic palmitoylation assay

Dynamic palmitoylation assay was performed as described previously^1^. In brief, OCI-AML3 cells were grown as described above, then spun down (3 minutes, 500*g*) and resuspended in fresh medium at a density of 2 × 10^6^ cells ml^−1^ (10 × 10^6^ cells per sample for gel-based assay and 20 × 10^6^ cells per sample for MS-based assay). Cells were preincubated with inhibitor or DMSO for 1 h, then 17-ODYA (20 µM) was added for 1 h. Samples were pelleted and snap frozen immediately following the 17-ODYA ‘pulse’ and designated *t*_0_, or resuspended in pre-warmed chase medium which consisted of OCI-AML3 growth media, supplemented with DMSO or inhibitor at the same concentration as in the preincubation step and incubated for 1 h. Cells were then centrifuged, placed on ice, washed in cold PBS, snap frozen and designated *t*_1_.

###### Immunoprecipitation

Cell pellets were resuspended in cold lysis buffer which consisted of 1% Triton-X PBS with 1 mM phenylmethyl sulfonyl fluoride (PMSF), 0.2 mM hexadecyl sulfonyl fluoride (HDSF), 20 µM HDFP and protease inhibitors (complete Ultra EDTA-free mini tablets, 5892791001 Roche), sonicated with a microtip probe sonicator (seven times, 50% rate, power 4), and placed on an end-over-end rotator for 30 minutes at 4°C. Samples were then hard spun (16,300*g*, 5 minutes) and the supernatant was transferred to fresh tubes on ice. Protein concentration was measured using a detergent compatible (DC) assay kit from Bio-Rad and adjusted to 1 mg ml^−1^. Input samples were taken at this point and stored at −80°C.

Enrichment was performed with anti-GFP Sepharose beads (20 µl of 50:50 slurry, ab69314, Abcam; RRID:AB_1640178). Antibody conjugated beads were spun down (500*g*), storage buffer was aspirated with a 26-gauge needle, and beads were washed three times with 1% Triton-X PBS. To each sample, 20 µl of washed bead slurry was added using a cut pipette tip, and samples were placed on an end-over-end rotator for 3 hours at 4°C. Next, washes were performed by centrifugation/aspiration (3×, 500*g*) with cold 1% Triton-X PBS containing 500 mM NaCl. After the last wash, supernatant was removed, and a 26-gauge needle was quickly inserted into the bead slurry to remove all remaining liquid. GFP beads were then resuspended in 50 µl of wash buffer. At this point, immunoprecipation samples were either stored at −80°C or processed further for analysis by SDS–PAGE gel.

###### Click chemistry and processing for SDS–PAGE gel

On-bead click chemistry was performed to conjugate a rhodamine fluorophore reporter to 17-ODYA labeled proteins. To each 50 µl sample, 6 µl of a click chemistry reaction mixture was added. The click reaction mixture was freshly prepared as follows (amounts given are per sample): 1 µl of 50 mM CuSO_4_ (in water; final concentration during reaction of 1 mM), 3 µl of 1.7 mM tris(benzyltriazolylmethyl)amine (4:1 *t*-BuOH/DMSO; 100 µM final), 1 µl of 50 mM TCEP (freshly made in PBS; 1 mM final) and 1 µl of Rh-N_3_ (DMSO; 25 µM final). Reactions were allowed to proceed for 1 hour at room temperature. Samples were then quenched with 4× SDS loading buffer containing 1% β-mercaptoethanol (described below), boiled for 5 minutes to effect elution and finally 17-ODYA proteins were resolved and imaged on gel.

Loading buffer was prepared as follows (for 100 ml): 3.02 g of Tris base was added to 40 ml of water, 40 ml of glycerol was added slowly and the mixture was stirred using a stir bar while the pH was brought to 6.75 using concentrated HCl; 8 g of SDS were then added followed by 20 ml of water and a pinch of bromophenol blue. Loading buffer was stored at room temperature, and 1% (10 µl per 1 ml of buffer) β-mercaptoethanol was added immediately before quenching.

Finally, either 15 µg protein samples were loaded on to a 4-20% Criterion^TM^ TGX gels (Bio-Rad) resolved by SDS–PAGE (160 V for ∼1 hour) or 30 µg protein samples were loaded on to a 10% acrylamide gel, resolved by SDS–PAGE (300 V for ∼2.5 hours), and in-gel fluorescence was visualized using a Bio-Rad ChemiDoc MP flatbed fluorescence scanner.

###### Cell viability analysis

Cell lines were plated at 1,000 cells/well on white opaque 96-well flat bottom plates (Perkin Elmer) in technical triplicate. Cells were treated with indicated drug dose series for 72 hours in culture conditions described in “Cell lines and tissue culture” section. MOLM-13 lab-generated lines were incubated in doxycycline-containing media (2mg / mL) for 24 hours prior to 96-well plating, and 72-hour experiments were carried out in the presence of 10nM quizartinib in addition to the indicated drug doses. Metabolically active cells as a proxy for overall cell viability were measured with CellTiter-Glo (Promega) and analyzed on a plate reader (Tecan M2000 Infinite Pro). Experiments were performed with technical triplicates and a representative experiment is shown from at least three experimental replicates for each cell line.

###### Cell cycle assay

OCI-AML3 cells (DSMZ, ACC-582) were plated at 50,000 cells/well on clear 96-well round bottom plates (Perkin Elmer) in technical triplicate. Cells were treated with the indicated drug doses for 24 hours in culture conditions described in “Cell lines and tissue culture” section. Plates were then removed from the incubator, centrifuged at 300 rcf for 5 minutes, and drug-containing media was removed by flicking. Cells were then fixed and permeabilized by resuspension in ice-cold 70% ethanol (diluted in ddH20) and stored at -20^0^ C for at least 30 minutes and up to 3 days. Plates were then centrifuged (900 rcf x 5 minutes), flicked, and washed, first with ice-cold PBS and then with HBSS (Gibco #14175-095) supplemented with 2% FBS. Cells were subsequently resuspended in HBSS + 2% FBS buffer containing 0.5ug/mL DAPI (BD #564097) and incubated for 15 minutes at room temperature. Staining media was removed by centrifugation and flicking, cells were resuspended in HBSS + 2% buffer and run on an Attune Nxt flow cytometer (ThermoFisher). DAPI^bright^ cells were considered to be dividing (“S + G2 + M”) based on the increased amount of DNA content in the cells.

###### Apoptosis assay

OCI-AML3 cells (DSMZ, ACC-582) were plated at 50,000 cells/well on clear 96-well round bottom plates (Perkin Elmer) in technical triplicate. Cells were treated with the indicated drug doses for 48 hours in culture conditions described in “Cell lines and tissue culture” section. Plates were then removed from the incubator, centrifuged at 300 rcf for 5 minutes, and drug-containing media was removed by flicking. Plates were then washed with HBSS (Gibco #14175-095) supplemented with 2% FBS, centrifuged, and flicked. Cell pellets were then resuspended in HBSS + 2% FBS buffer containing 80nM ApoTracker Green (BioLegend #427402) and DAPI 0.2ug/mL (BD #564097) and incubated for 15 minutes in the dark at room temperature. Staining media was then removed by centrifugation and flicking, and the plates were washed again as described above. ApoTracker and DAPI-stained cells were then resuspended in HBSS + 2% FBS and run on an Attune NxT flow cytometer (ThermoFisher). DAPI^+^ / ApoTracker^+^ cells were considered “dead”, DAPI^-^/ ApoTracker^+^ cells were considered “apoptotic”.

###### Colony forming assay

Incomplete MethoCult^TM^ (STEMCELL Technologies) was supplemented with IMDM +/- GM-CSF and ABD778 (final concentration 1μM) or vehicle per the manufacturer’s recommended protocol. Cryopreserved patient JMML bone marrow cells were mixed with supplemented MethoCult^TM^ to achieve a final concentration of 2750 CD34^+^ cells / 3.3mL media. 1.2mL of cell / MethoCult^TM^ mixture (containing 1000 CD34^+^ cells) were aliquoted in duplicate into 30mm plates, and then incubated for 14 days at 37 °C with 5% CO2 prior to colony counting.

###### Phospho-flow cytometry

Cells were incubated with drug and/or vehicle for 4 or 24 hours in culture conditions described in “Cell lines and tissue culture” section, then were fixed with a final concentration of 2% paraformaldehyde (Electron Microscopy Sciences) added directly to drug-containing media. For MOLM-13 cytokine stimulation experiments, the same procedure was followed except that cells were incubated in serum-free media with 10nM quizartinib and 2mg/mL doxycycline (all media components otherwise identical) for 24 hours, followed by addition of 100 ng/mL human IL-3 (PeproTech) for 10 minutes prior to fixation. Samples were washed as described above, permeabilized with ice-cold methanol and stored at -20^0^ C for at least 30 minutes and up to 3 days, washed again, then stained with pERK (Cell Signaling Cat#9101, RRID:AB_331646), pS6 (Cell Signaling Cat#4858, RRID:AB_916156), or pSTAT5 (Cell Signaling Cat#9359, RRID:AB_823649). Cells were then washed, stained with a secondary antibody (Jackson Immunoresearch Cat#711-135-152, RRID:AB_2340601), and analyzed on a LSR II flow cytometer (BD Biosciences).

###### Immunoblotting

Cells were incubated with drug and/or vehicle for 4 or 24 hours, then were lysed in ice cold RIPA buffer (Pierce) supplemented with HaltTM protease and phosphatase inhibitor cocktail (Thermo Fisher) and 0.5 µM EDTA, incubated on ice for 30 minutes, and centrifuged at maximum speed for 15 minutes to collect whole cell lysates. Protein concentration was measured with the BCA protein assay (Pierce). 30µg of total protein per sample was loaded into 4-12% gradient Criterion^TM^ TGXTM gels (Bio-Rad) and separated by SDS-PAGE. Proteins were transferred to PVDF or nitrocellulose membranes and blotted with the following primary antibodies: pERK (Cell Signaling Cat#9101, RRID:AB_331646), ERK (Cell Signaling Cat#9107, RRID:AB_10695739), pS6 (Cell Signaling Cat#4858, RRID:AB_916156), S6 (Cell Signaling Cat#2317, RRID:AB_2238583), pMEK (Cell Signaling Cat#2338, RRID:AB_490903), MEK (Cell Signaling Cat#4694, RRID:AB_10695868), pAKT (Cell Signaling Cat#4060, RRID:AB_2315049), AKT (Cell Signaling Cat#2920, RRID:AB_1147620). GAPDH (Cell Signaling Cat#97166, RRID:AB_2756824), Ras^G12D^ (Cell Signaling Cat#14429, RRID:AB_2728748), and Hsp90 (Cell Signaling Cat#4875, RRID:AB_2233331) was used for loading controls.

###### NanoBRET Assay

NanoBRET assays were performed as described by Promega. HEK293T cells (ATCC Cat# CRL-3216) were cultured at 37°C, 5% CO2 in DMEM, high glucose (Gibco Cat# 11965092) supplemented with 10% FBS, 1% penicillin/streptomycin, and 1% non-essential amino acids. Cells were transfected with HaloTag (HT)-NRAS (WT or Q61L) and RAF-NanoLuciferase (Nluc) (A-, B-, or C-RAF) in a 1:10 ratio of RAF-Nluc:HT-NRAS plasmid using jetOPTIMUS transfection reagent (Sartorius Cat# 101000006). After 24 hours, cells were reseeded at 8,000 cells per well in 384-well plates (Corning Cat# 3570) in FluoroBrite DMEM (phenol red-free; Gibco Cat#A1896701) with 4% FBS and 1% penicillin/streptomycin. Cells were treated with 100 nM HaloTag NanoBRET 618 Ligand (Promega Cat# G9801) or same % v/v DMSO as a “no ligand negative control”. The extended live cell substrate, Nano-Glo Endurazine substrate (Promega Cat# N2571), was added to the cells to allow for continual reading of the assay plates. ADB778 and the control RMC-7977 (MedChemExpress Cat# HY-156498) were tested in a 2-fold dilution series starting at 3 μM (ADB778) or 1 μM (RMC-7977) with a final % DMSO of 0.25%. Assay plates were read after a 1-hour incubation using an EnVision Xcite 2105 plate reader 618 nM (HaloTag) and 460 (Nluc). The final corrected NanoBRET ratio was calculated using the following equation:

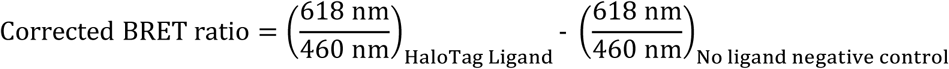

Data was visualized using GraphPad Prism 10, using log(agonist) vs. response – variable slope (four parameters) nonlinear regression curve fit.

###### *In vivo* pharmacodynamic analyses of ABD778 and PD901

8-12 week old F1 mice (first generation hybrid of B6 x 129/Sv mice) were dosed daily with ABD778 (60 mg/kg), PD901 (2.5 mg/kg), this combination, or vehicle by oral gavage for 4 days. At either 2, 6, or 24 hours after the final dose of drug, mice were sacrificed and bone marrow cells were collected from one hindlimb. Homogenized and filtered bone marrow cells underwent RBC lysis, then were washed and plated in starve media (IMDM with 1% penicillin/streptomycin, 2mM L-glutamine, and 1% BSA) at 500,000 cells / well in a 96-well round bottom plate (Perkins Elmer). Cells were then stimulated with 10ng/mL murine GM-CSF (PeproTech) for 10 minutes, and were subsequently fixed, permeabilized, stained, and analyzed as described in the “Phospho-flow cytometry” section. Antibodies used for staining were against pERK (Cell Signaling Cat#4370, RRID:AB_2315112), Rabbit IgG (Jackson ImmunoResearch Cat#111-117-008, RRID:AB_2632461), CD11b / Mac1 (BD Biosciences Cat#561690, RRID:AB_10897015), and pSTAT5 (BD Biosciences Cat#560311, RRID:AB_1645497).

###### Preclinical trials in mouse AMLs

Transplantable primary AMLs were generated by injecting neonatal wild-type (WT), *Mx1-Cre; Nras^LSL-G12D/+,^* or *Mx1-Cre; Kras^LSL-G12D/+,^*mutant mice with the MOL4070LTR retrovirus followed by a single dose of polyI-polyC at weaning to induce *Nras^G12D^*or *Kras^G12D^* expression from the endogenous loci as described (see Refs 37, 42). Preclinical trials were performed by expanding cryopreserved leukemia cells *in vivo* and transplanting 1 x10^6^ bone marrow cells into irradiated (450cGy) 8-to 12-week-old congenic recipients. Mice were dosed daily with ABD778 (60 mg/kg), PD901 (2.5 mg/kg), this combination, or vehicle by oral gavage. Drug and vehicle were made fresh each week. Mice were monitored daily until they became symptomatic, then twice daily until moribund. Bone marrow, spleen, and cardiac blood was collected for genomic DNA extraction, cryopreservation, and complete blood count analysis (HemaTrue).

###### Whole exome sequencing

Genomic DNA was extracted from the bone marrow cells and then sheared to generate 150 to 200 base pair fragments using a Covaris S2 focused-ultrasonicator. Indexed libraries were prepared using the Agilent SureSelect XT2 Reagent Kit for the HiSeq platform. Exomes were captured using the Agilent SureSelect XT2 Mouse All Exon bait library. Sample quality and quantity were assessed using the Agilent 2100 Bioanalyzer instrument. Paired-end 100 base pair reads were generated on an Illumina HiSeq 2000 platform. All sequence data including read alignment; quality and performance metrics; post-processing, somatic mutation and DNA copy number alteration detection; and variant annotation were performed as previously described^9^ using the mm10 build of the mouse genome. Briefly, reads were aligned with Burrows-Wheeler Aligner, and processed using Picard (http://broadinstitute.github.io/picard) tools and the Genome Analysis Toolkit (GATK) to perform base quality recalibration and multiple sequence realignment. Single nucleotide variants and indels were detected with the MuTect and Pindel algorithms, respectively. Candidate somatic mutations were manually reviewed using Integrative Genomics Viewer.

###### Plasmids, cloning, and mutagenesis

The plasmids pCW57.1 (Addgene plasmid 41393) and pTwist-ENTR BRAF^G466E^-T2A-mCherry (custom order, Twist Biosciences) were used to generate a dox-inducible pCW57.1-BRAF^G466E^-mCherry construct with Gateway LR Clonase enzyme mix (Thermo Fisher). The dox-inducible pCW57.1-mCherry-Kras^A146T^ vector had been previously generated by similar methods (see Ref 31).

###### Lentiviral generation and transduction

Lentiviral backbone, pMD2.G (Addgene plasmid #12259) and psPAX2 (Addgene plasmid #12260) were transfected into 293T lenti-X cells (Takara Bio) with TransIT-LT1 (Mirus Bio). Supernatant was collected 48 hours after transfection and applied to OCI-AML3 cells with polybrene for transduction. Cells were spin-infected at 800*g* for 2 hours at 37°C. After recover, cells were treated with 2 μg/mL doxycycline (Sigma) and transduction efficiency was determined by BD LSR II analysis. MOLM13 lines were previously generated by similar methods (see Ref 31).

###### Competitive growth assay

Transduced OCI-AML3 cells were grown in the presence of 2 µg/mL doxycycline and drug or vehicle as indicated. Cell viability (Beckman Coulter Vi-Cell XR) and fluorescent protein expression (BD LSRII) were measured every 2 days for 10 total days. After measurements, cells were re-plated with fresh doxycycline and drug or vehicle at a density of 0.5 x 10^6^ cells / well.

###### Melanoma growth assays and immunoblotting

D04, MM415, MM485, and SK-MEL-2 cells were cultured in drug-containing media at the indicated doses for 3-5 days before analysis as described in the “Cell viability assay” section. Similarly, melanoma lines were exposed to drug-containing media for 4, 24, or 48 hours before lysis and immunoblotting as described in the “Immunoblotting” section.

###### Patient-derived xenograft trial

NOD.Cg-Prkdc^scid^ Il2rg^tm1Wjl^/SzJ (NSG) were obtained from in-house breeding stocks at the UCSF Preclinical Therapeutics Core facility housed and bred in a pathogen-free facility. A mixture of male and female mice between 6 and 12 weeks old were transplanted with 1e6 viably frozen passaged PDX spleen cells via tail vein injection after two consecutive days of busulfan conditioning (25 mg/kg) and one day of rest. Tumor engraftment was assessed by flow cytometry for hCD45^+^ percentage in PB from submandibular bleeding on day 4. Mice were randomized to receive one of four treatments, six mice per treatment arm: (1) ABD778 50 mg/kg daily, (2) gilteritinib 30 mg/kg daily (C) ABD778 50 mg/kg daily and gilteritinib 30 mg/kg daily or (4) vehicle. Mice were analyzed for hCD45^+^ at the mid-point of the trial via submandibular bleeding. All mice were sacrificed 21 days after initiation of treatment at which point cardiac blood, spleen, and bone marrow cells were analyzed for hCD45^+^.

###### Statistical Analysis

Statistical analyses were performed using GraphPad Prism 10. Multiple comparisons between treatment conditions were performed using one way ANOVA and *Šídák* multiple comparisons testing. Kaplan Meier statistics were calculated by two-sided log rank test. In the figures, the degree of significance is denoted by the number of asterisks (\*\*\*\**P* < 0.0001, \*\*\**P* ≥ 0.0001 and < 0.001, \*\**P* ≥ 0.001 and < 0.01, \**P* ≥ 0.01 and < 0.05, NS ≥ 0.05). Error bars indicate the mean ± s.d. unless otherwise indicated in the figure legend.

## Notes

### Summary of Updates

Figure 2 and 3 revised to include new mechanistic data. Previous Figure 4 and 5 (in vivo murine trial data) combined. New Figure 5 with melanoma cell line data. Figure 6 updated to include RAS(ON) inhibitor synergy. Text revised to describe new findings. Author list updated to reflect new contributors.

