## Supplementary Figures and Methods for "*In Vivo* Efficacy of ABHD17 Inhibition Validates the Palmitoylation Cycle as a Therapeutic Target in *NRAS*-Mutant Cancer"


(1) Figures S1-15

(2) Tables S1-4

(3) Extended Methods: Synthesis and Characterization of ABD778

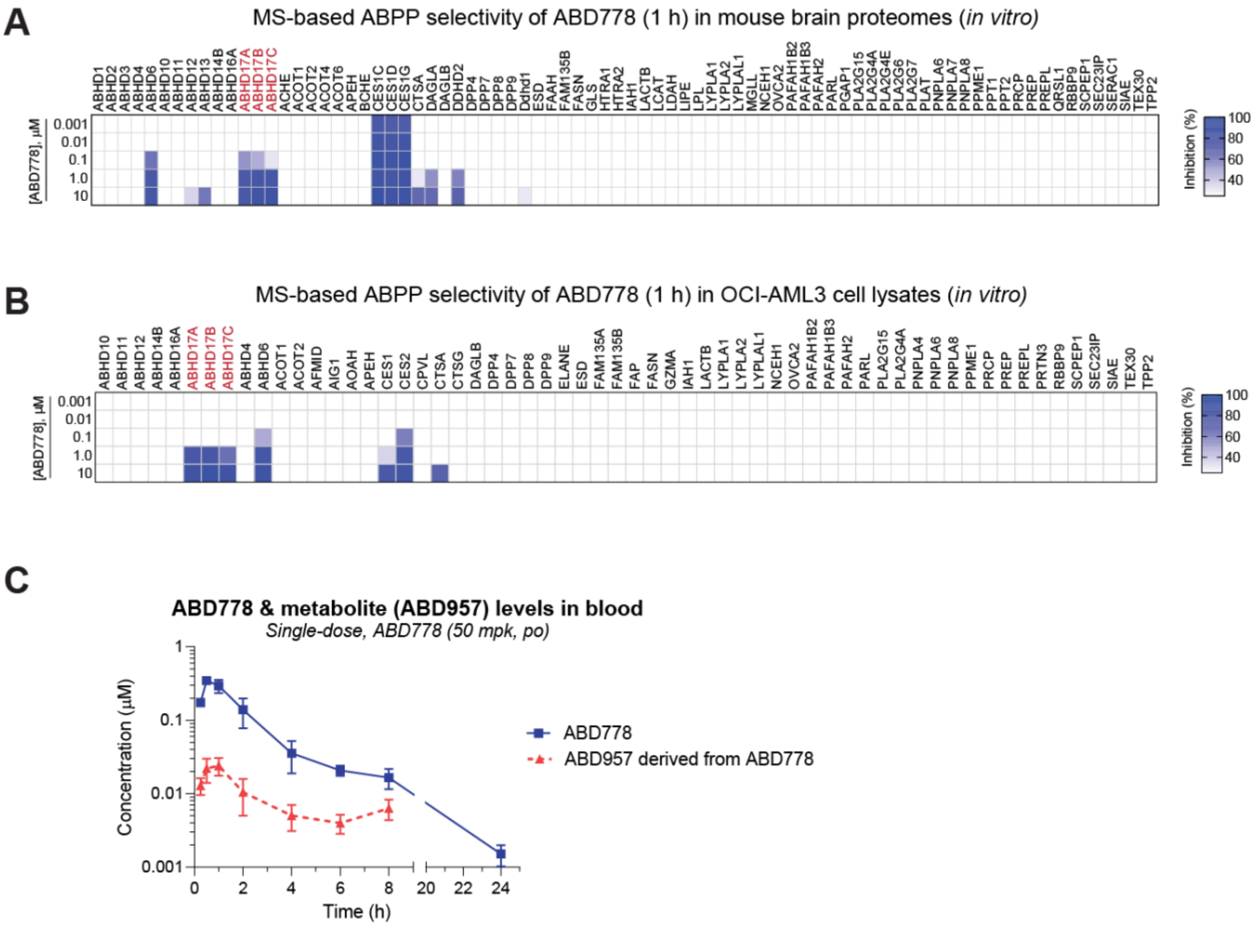


**Figure S1. Characterization of ABD778 as an ABHD17 inhibitor with *in vivo* activity. (A)(B)** In depth serine hydrolase selectivity profiling by MS-based ABPP for ABD778 (0.001–10 µM, *in vitro*) in mouse brain **(A)** and OCI-AML3 **(B)** cell proteomes confirming ABHD17A/B/C inhibition and selectivity across the serine hydrolome including for the depalmitoylases LYPLA1, LYPLA2 and ABHD10. Prominent off-targets include ABHD6 and carboxyesterases (CESs). Data plotted represent mean competition from three biological replicates. **(C)** Blood concentrations of ABD778 and ABD957 following a single dose of ABD778 (50 mg/kg, po) in mice. Blood was serially collected at the indicated timepoints following compound administration and analyzed LC-MS/MS. Note that the oncentrations of ABD957 arising from the esterolysis of ABD778 (red dotted line) are far below those of ABD778 (blue line).

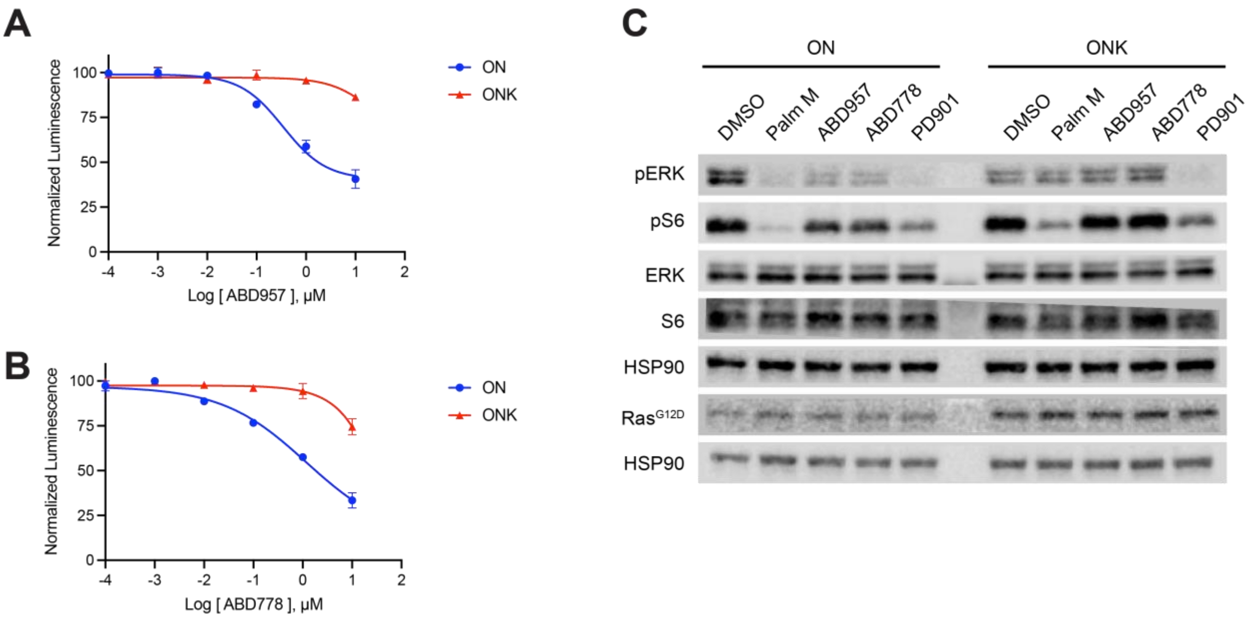


**Figure S2. (A)(B)** Cell growth as assessed by Cell Titer Glo in OCI-AML3/ON (ON) and OCI-AML3/ONK (ONK) cells exposed to a range of **(A)** ABD957 or **(B)** ABD778 doses. **(C)** Western blot analysis of ON and ONK cells were exposed to either the promiscuous serine hydrolase inhibitor Palmostatin M (10 μM), ABD957 (500 nM), ABD778 (500 nM), or PD901 (10 nM) for 3 hours showing levels of total (t) and phosphorylated (p) ERK1/2 and S6. Antibodies to HSP90 and $Ras^{G12D}$ were used as loading controls.

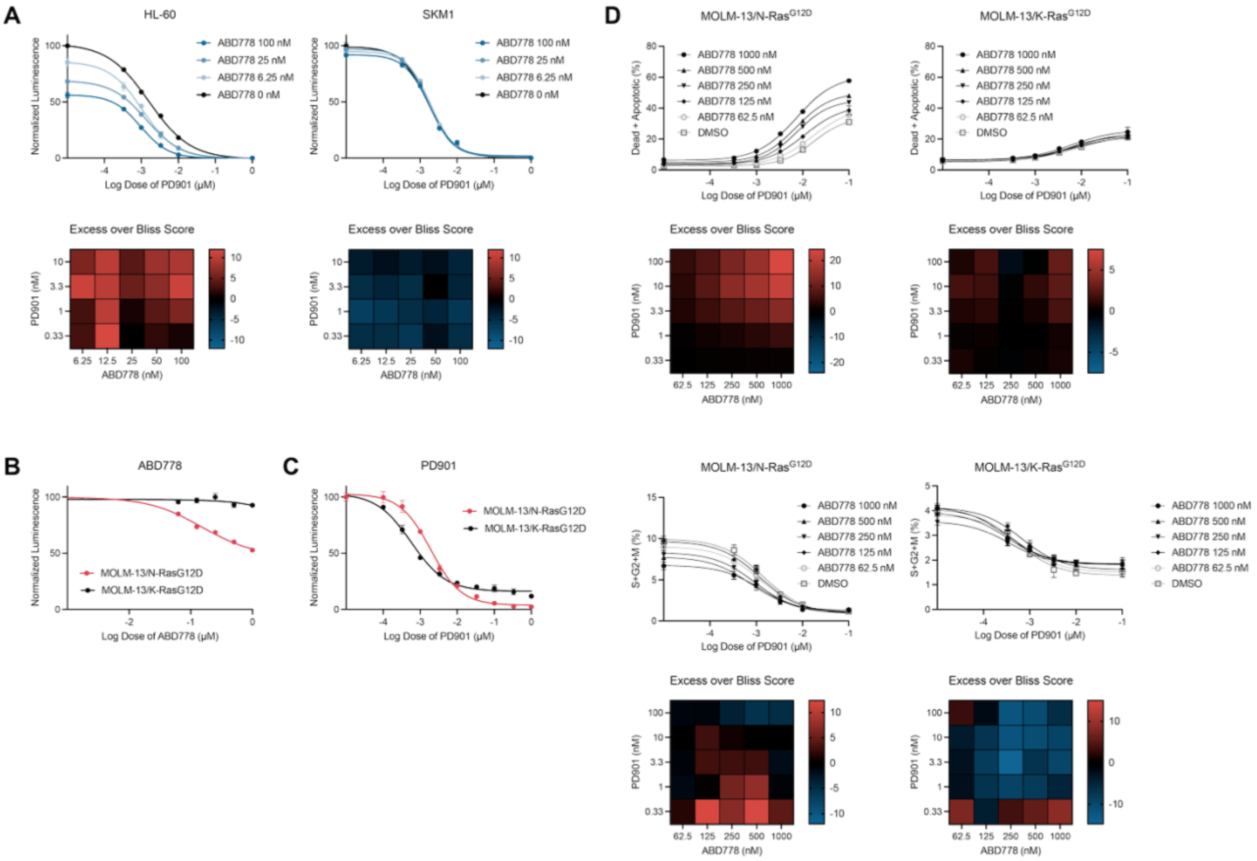

**Figure S3. Selective activity of ABD778 and synergy with PD901 in *NRAS*-mutant AML cell lines. (A)** Bliss independence analysis demonstrating synergistic growth inhibition over a broad range of ABD778 and PD901 concentrations in *NRAS*-mutant HL-60 (left), but not *KRAS*-mutant SKM1 (right), AML cells. **(B, C)** MOLM-13 cells that were treated with doxycycline (Dox) for 24 hours to induce exogenous N-Ras$^{G12D}$ (MOLM-13/N-Ras$^{G12D}$) or K-Ras$^{G12D}$ (MOLM-13/K-Ras$^{G12D}$) expression as described previously[31,32] were exposed a range of ABD778 and PD901 doses for 72 hours. Proliferation was assessed by Cell Titer-Glo. **(D)** Bliss independence analysis of apoptosis and cell cycling in MOLM-13/N-Ras$^{G12D}$ (left) and MOLM-13/K-Ras$^{G12D}$ (right) cells treated with ABD778 and PD901. Heatmaps in panels A and D display calculated synergy scores from strongly positive (red) to negative (blue). These data were generated in triplicate and were replicated in at least two additional independent experiments.

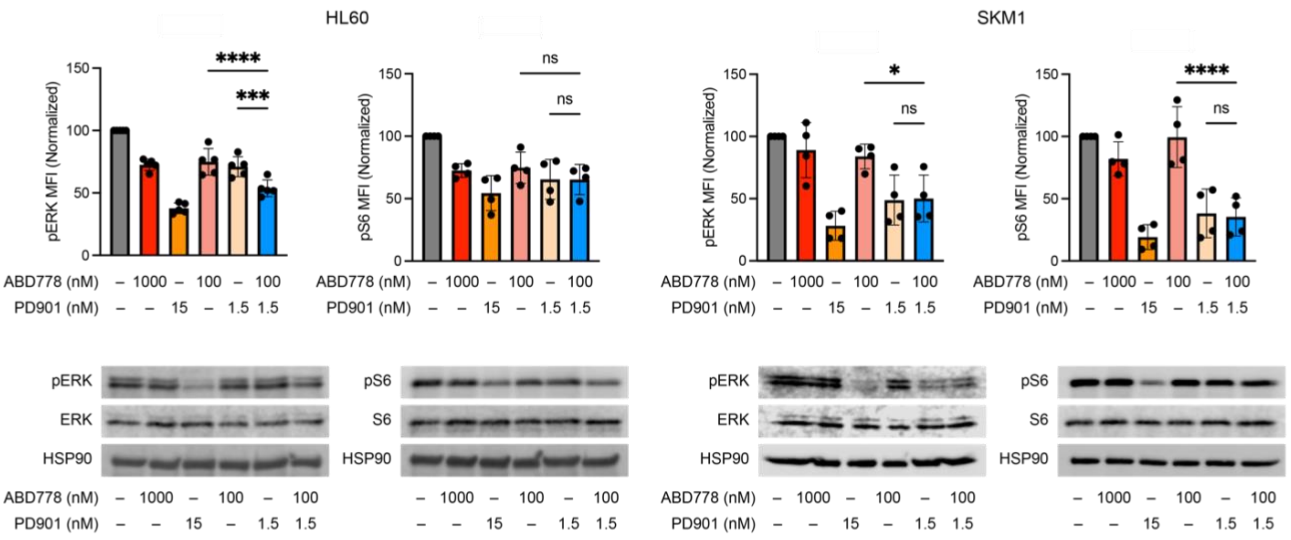


**Figure S4. Biochemical analysis of HL-60 and SKM1 AML cells that were exposed to DMSO vehicle, ABD770, PD901, or both drugs.** Top**.** Phosphorylated ERK **(**pERK) and S6 (pS6) levels were measured using phospho-flow cytometry in HL-60 (left) and SKM1 (right) cells that were exposed to ABD778 and/or PD901 for 4 hours at the doses shown. Mean fluorescence index (MFI) values were normalized to 100% of the DMSO control and pooled for statistical analysis (n = 4-5; * - $p < 0.05$; *** - $p < 0.001$; **** - $p < 0.0001$). Bottom**.** pERK and pS6 levels were measured by Western blotting in HL-60 (left) and SKM1 (right) cells that were exposed to DMSO, ABD778, PD901, or both drugs. The phospho-flow data were generated in three independent experiments and the Western blot data are respentative of at least three independent experiments.

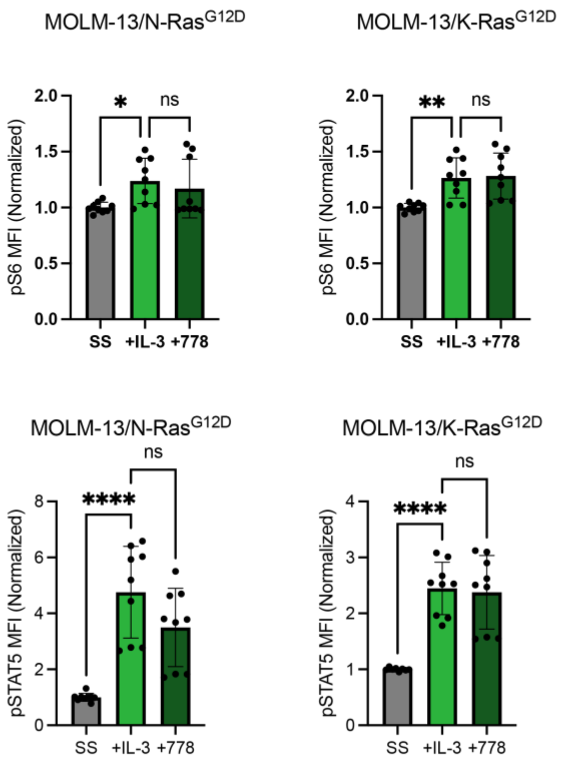


**Figure S5. ABD778 does not reduce S6 or STAT5 phosphorylation in response to IL-3 stimulation.** pS6 (top) and pSTAT5 (bottom) levels were measured using phospho-flow cytometry in MOLM-13/N-Ras$^{G12D}$ (left) and MOLM-13/K-Ras$^{G12D}$ (right) cells after 24 hours in serum-free media + DMSO (SS), 24 hours in serum-free media + DMSO followed by IL-3 stimulation (+IL-3), and 24 hours in serum-free media + 100nM ABD778 followed by IL-3 stimulation (+778). n = 9; * - $p < 0.05$; ** - $p < 0.01$; **** - $p < 0.0001$

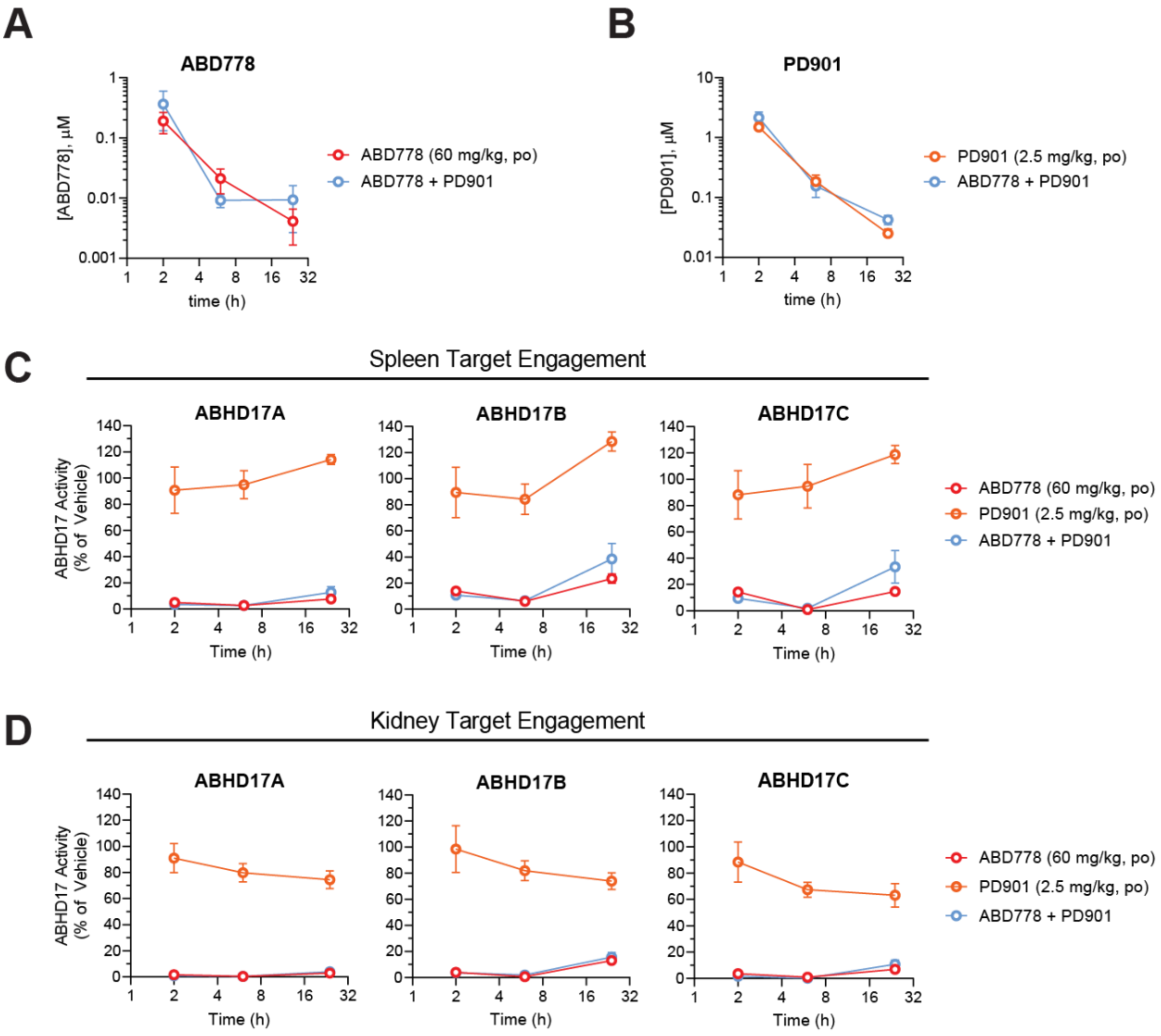


**Figure S6. Single agent and combination treatment result in similar drug levels and inhibition of ABHD17 activity in hematopoietic tissues.** Blood concentrations of ABD778 (**A**) and PD901 (**B**) following a single dose of ABD778 (60 mg/kg, po), PD901 (2.5 mg/kg, po) or both compounds dosed 1 hour apart in mice. Blood was collected 2, 6, and 24 hours after compound administration and analyzed by LC-MS/MS. Co-administration of ABD778 and PD901 (blue line) provides similar exposures compared with single-agent dosing (red lines). **C**, **D**, *In vivo* target engagement for ABHD17A/B/C in spleen (**C**) and kidney (**D**) measured by MS-based ABPP. C57Bl/6 mice were dosed with vehicle, ABD957 (60 mg/kg, po), PD901 (2.5 mg/kg, po) or both compounds dosed 1 hour apart. Tissues were collected 2, 6, and 24 hours after compound administration and analyzed by targeted MS-ABPP using FP-biotin enrichment of serine hydrolase enzymes and parallel reaction monitoring (PRM) to detect and quantify unique diagnostic peptides from ABHD17A, B and C. ABD778 provided near complete blockade of ABHD17 enzymes in spleen and kidney across the 24 hour time course when dosed as a single agent or in combination with PD901. Data plotted represent the median from biological replicates, and error bars represent s.e.m. (n = 3-4).

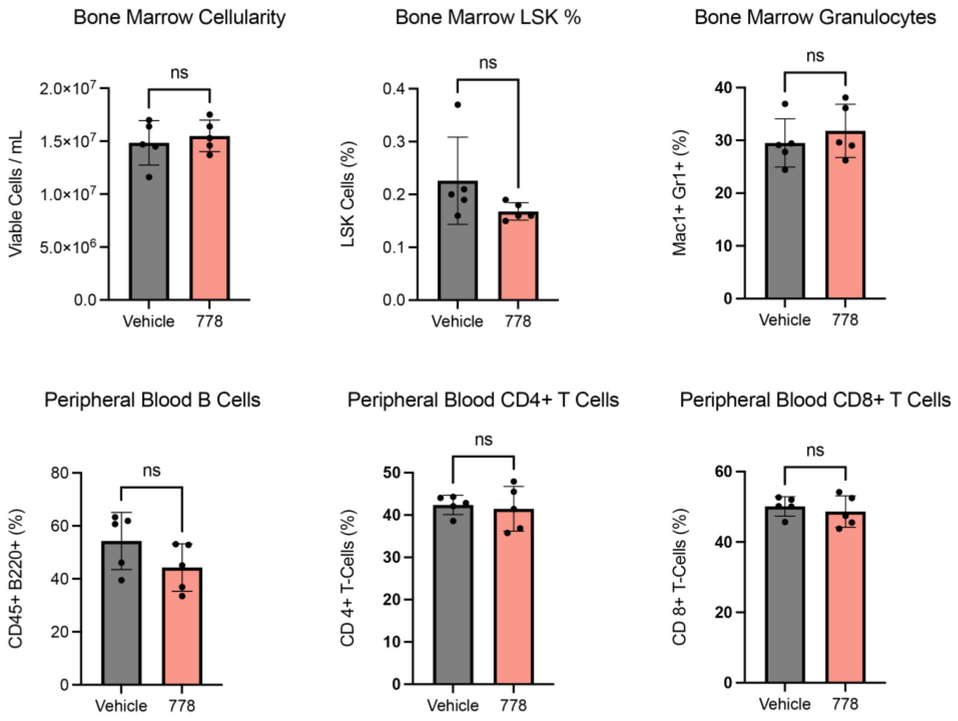


**Figure S7. ABD778 treatment does not acutely alter blood and bone marrow populations in mice.** Wild-type mice were treated with ABD778 (60mg/kg, po) or vehicle for 4 days then euthanized. Peripheral blood and bone marrow was collected, processed, and immunostained for stem and progenitor cells (LSK%), innate immune cells (granulocytes), and adaptive immune cells (B-cells, CD4+ T-cells, CD8+ T-cells) as labeled.

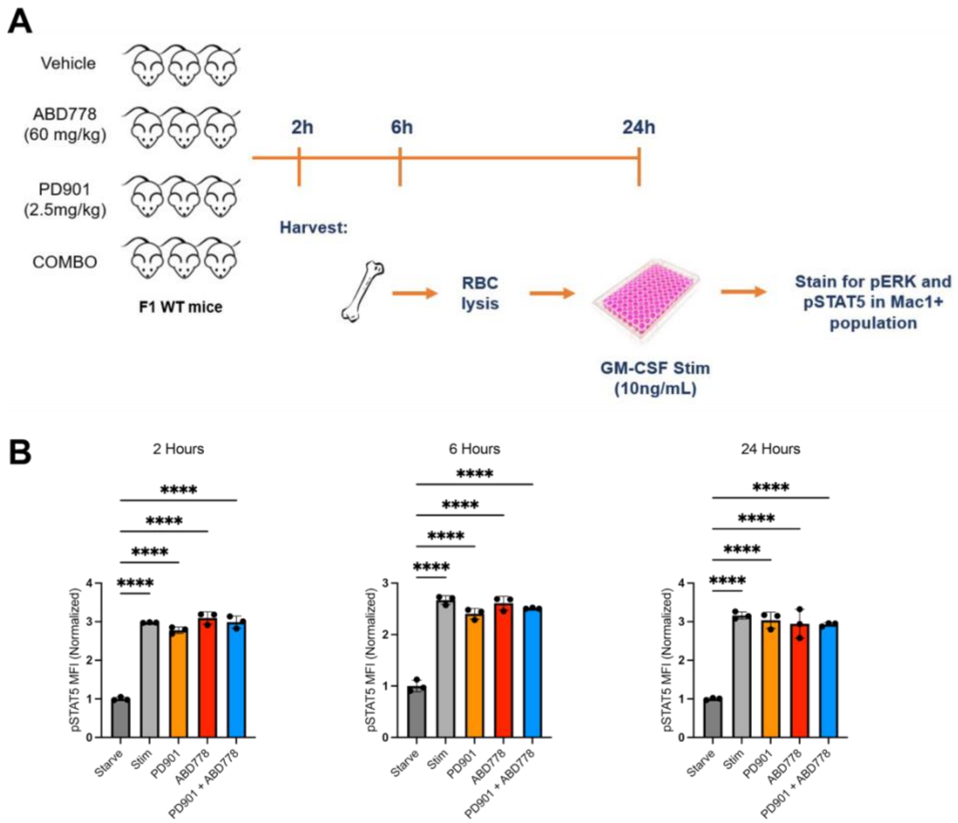


**Figure S8. GM-CSF stimulation of bone marrow cells from wild-type might collected from mice after treatment with ABD778, PD901, or both compounds. (A**) Schema of assay. Briefly, mice were treated with vehicle, ABD778 (60mg/kg, po), PD901 (2.5mg/kg, po), or both compounds for 4 days. Bone marrow cells were collected at 2, 6, or 24 hours after final dose administration, plated, and stimulated with 10ng/mL of GM-CSF. **(B)** Phospho-flow analysis of STAT5 activation in GM-CSF stimulated Mac1+ bone marrow cells. MFI normalized to unstimulated (Starve) condition at each timepoint. **** - $p < 0.0001$.

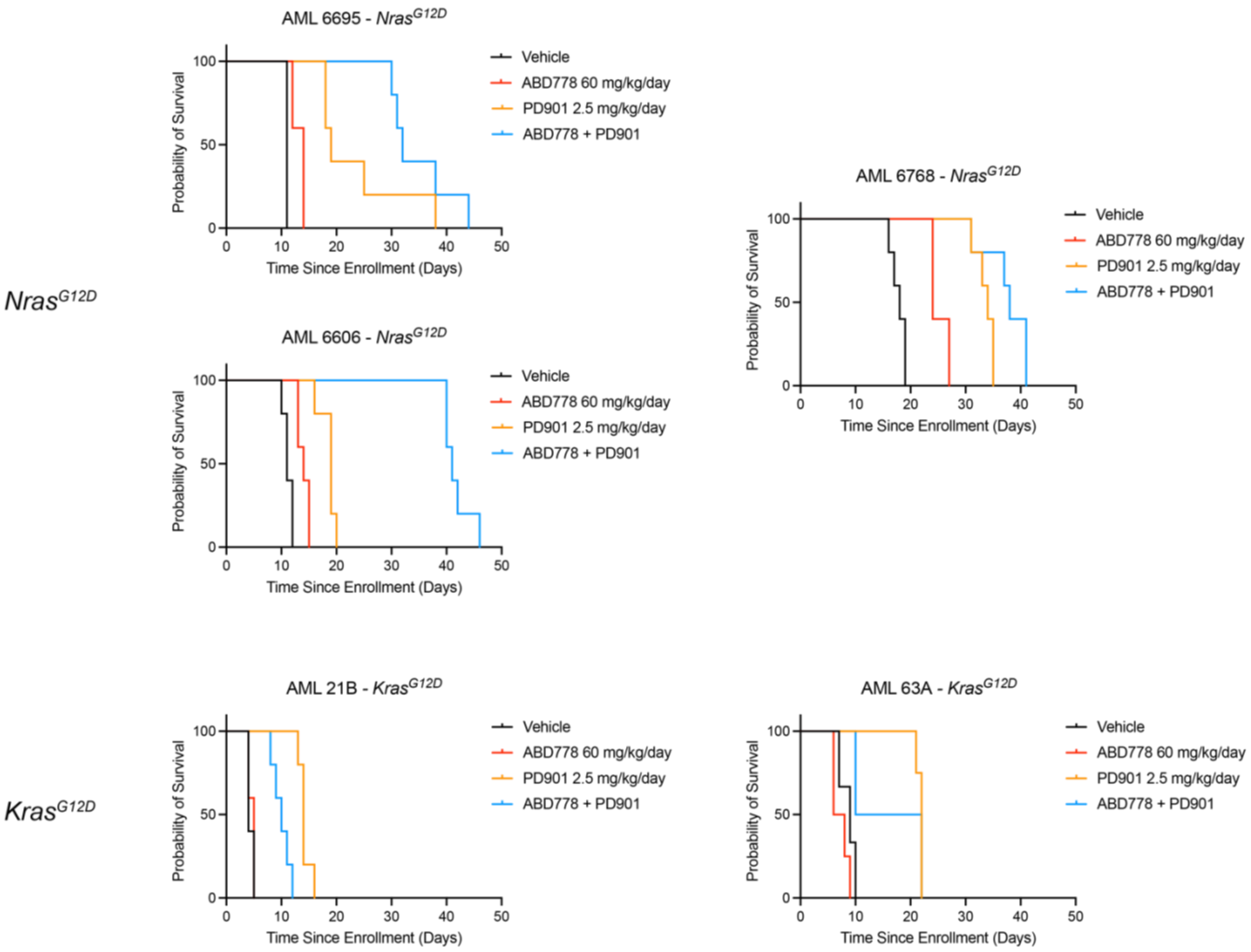


**Figure S9.** Kaplan-Meier survival analysis of primary *Nras*$^{G12D}$ (n = 3) and *Kras*$^{G12D}$ (n = 2) AMLs enrolled on preclinical trials of ABD778 and PD901 (n = 5 mice per group x 4 groups = 20 mice in each trial).

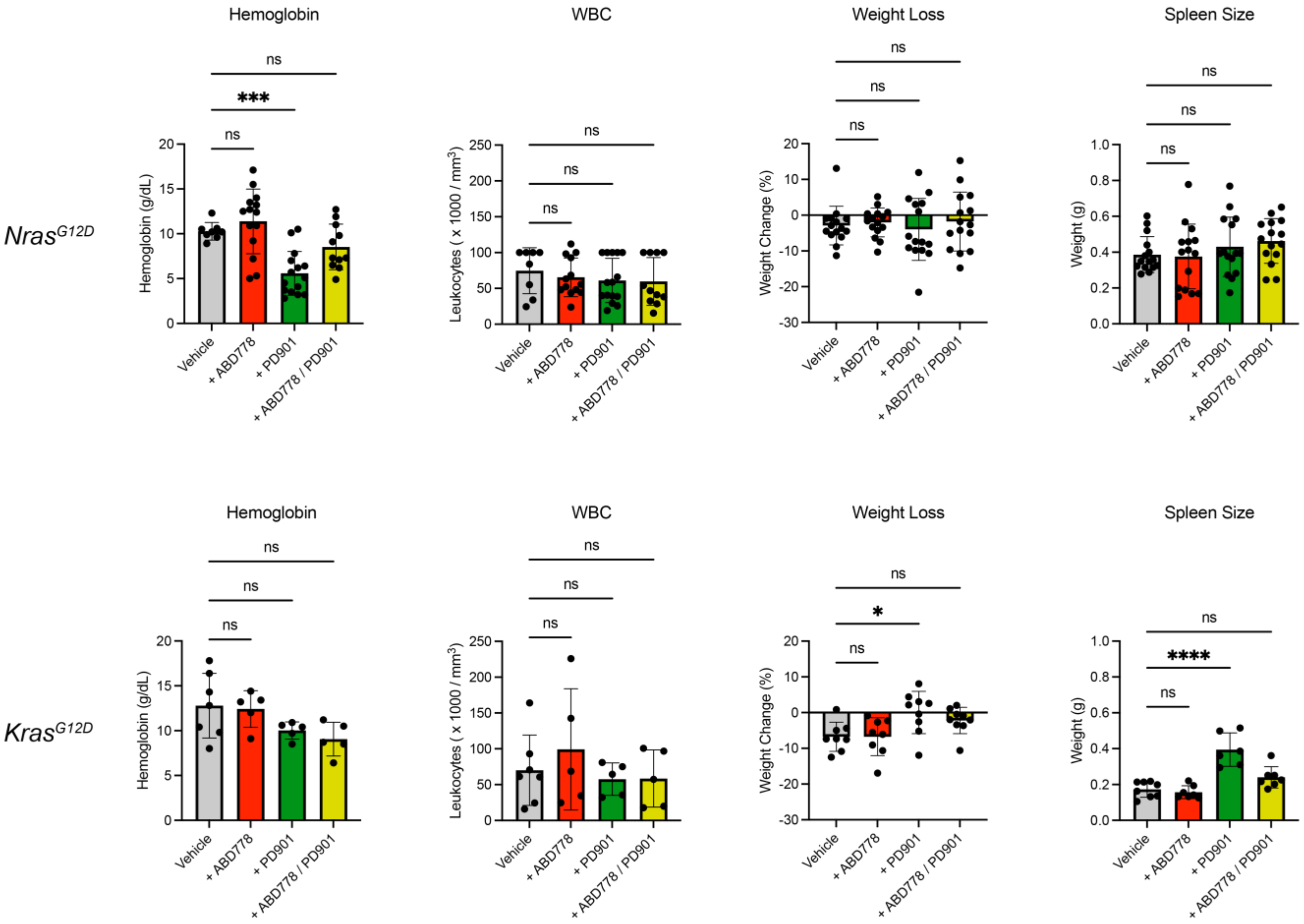


**Figure S10. Treatment-refractory AML in recipient mice at euthanasia.** Despite significant differences in overall survival in some treatment arms, recipient mice in all treatment groups died with anemia (normal > 13g/dL), elevated blood leukocyte counts (normal < 14,000 per $mm^3$), and splenomegaly (normal < 100mg), which is indicative of refractory AML. n = 5-15; * - p < 0.05; *** - p < 0.001; **** - p < 0.0001.

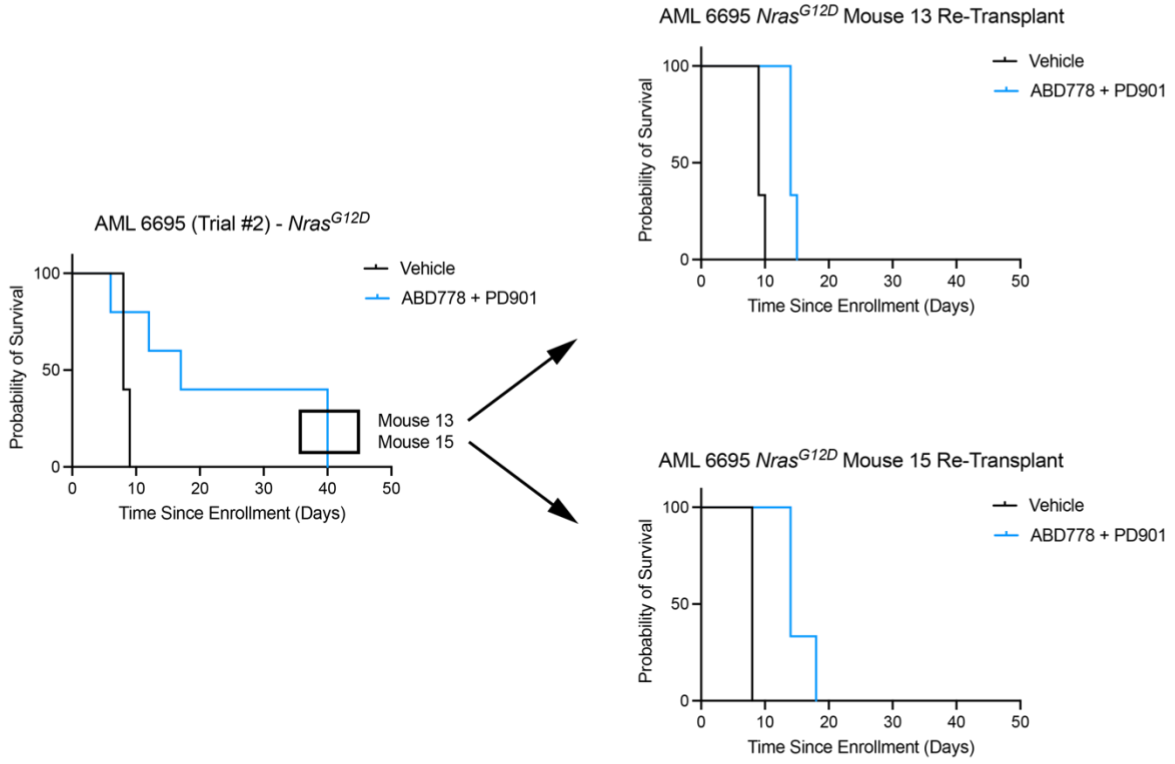


**Figure S11. Refractory AMLs that emerge after combination treatment exhibit intrinsic drug resistance.** The left panel shows the survival of 5 recipient mice that were transplanted with *Nras*$^{G12D}$ AML 6606 and treated ABD778 + PD901 (blue line; n =5). Leukemia cells collected at euthanasia from the two recipients shown in the black box were indpendently injected into secondary recipients and re-treated with either control vehicle or ABD778 + PD901 (n = 3 mice per group). The right panel shows the survival of these mice.

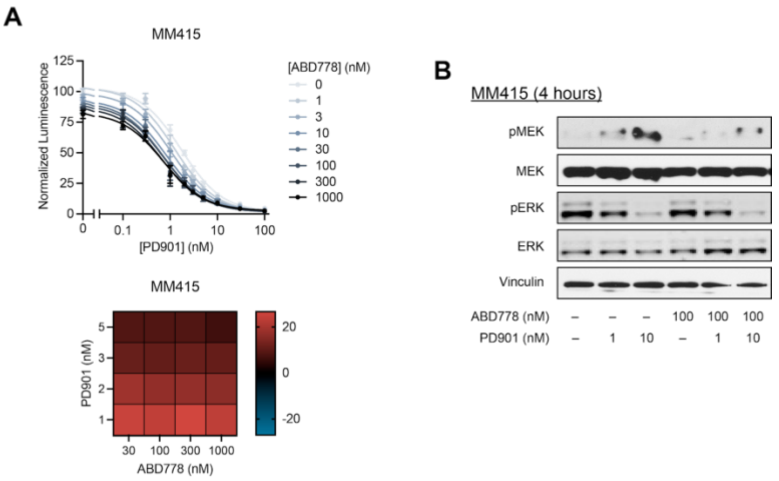


**Figure S12. PD901/ABD778 combination treatment synergistically blocks growth and decreases pERK levels in MM415 melanoma cells. A,** Proliferation of *NRAS*-mutant M415 melanoma cells was measured after exposure to ABD778 and PD901 using Cell Titer-Glo. Bliss independence analysis is displayed as a heatmap of calculated synergy scores from strongly positive (red) to negative (blue). **B.** pERK and pMEK levels were measured by Western blotting in MM415 cells that were exposed to DMSO, ABD778, PD901, or both drugs for 4 hours.

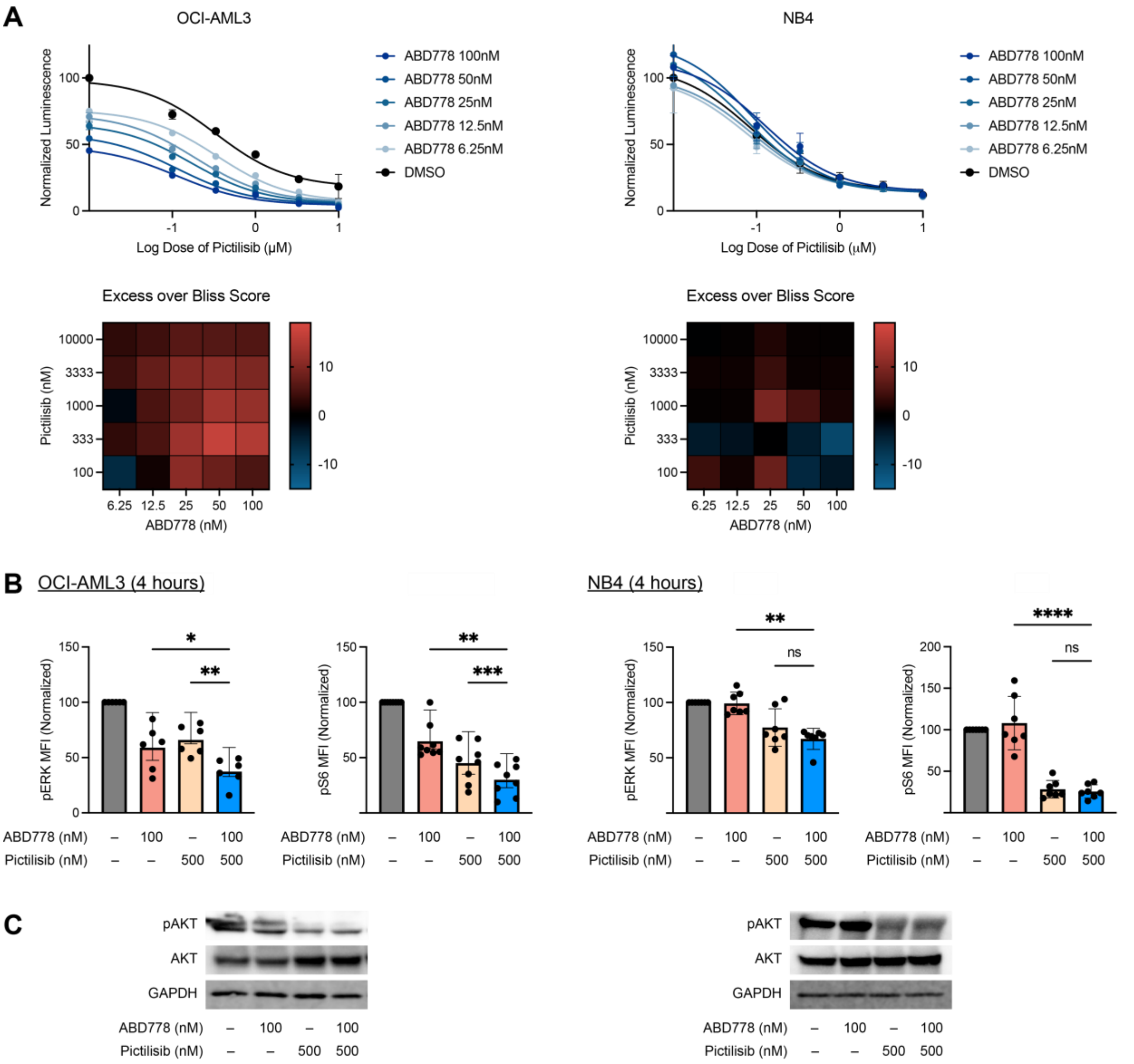


**Figure S13. Synergistic and biochemical analysis of OCI-AML3 and NB4 AML cells that were exposed to DMSO vehicle, ABD770, pictilisib, or both drugs. (A)** Bliss independence analysis demonstrating synergistic growth inhibition over a broad range of ABD778 and pictilisib concentrations in *NRAS*-mutant OCI-AML3 (left), but not *KRAS*-mutant NB4 (right), AML cells. Heatmaps display calculated synergy scores from strongly positive (red) to negative (blue). These data were generated in triplicate and were replicated in at least two additional independent experiments. **(B)** Phosphorylated ERK (pERK) and S6 (pS6) levels were measured using phospho-flow cytometry in OCI-AML3 (left) and NB4 (right) cells that were exposed to ABD778 and/or pictilisib for 4 hours at the doses shown. Mean fluorescence index (MFI) values were normalized to 100% of the DMSO control and pooled for statistical analysis (n = 6-8; **** - p < 0.0001). **(C)** pAKT levels were measured by Western blotting in OCI-AML3 (left) and NB4 (right) cells that were exposed to DMSO, ABD778, PD901, or both drugs. The data presented were generated in (phospho-flow) or are representative of (Western blots) at least three independent experiments.

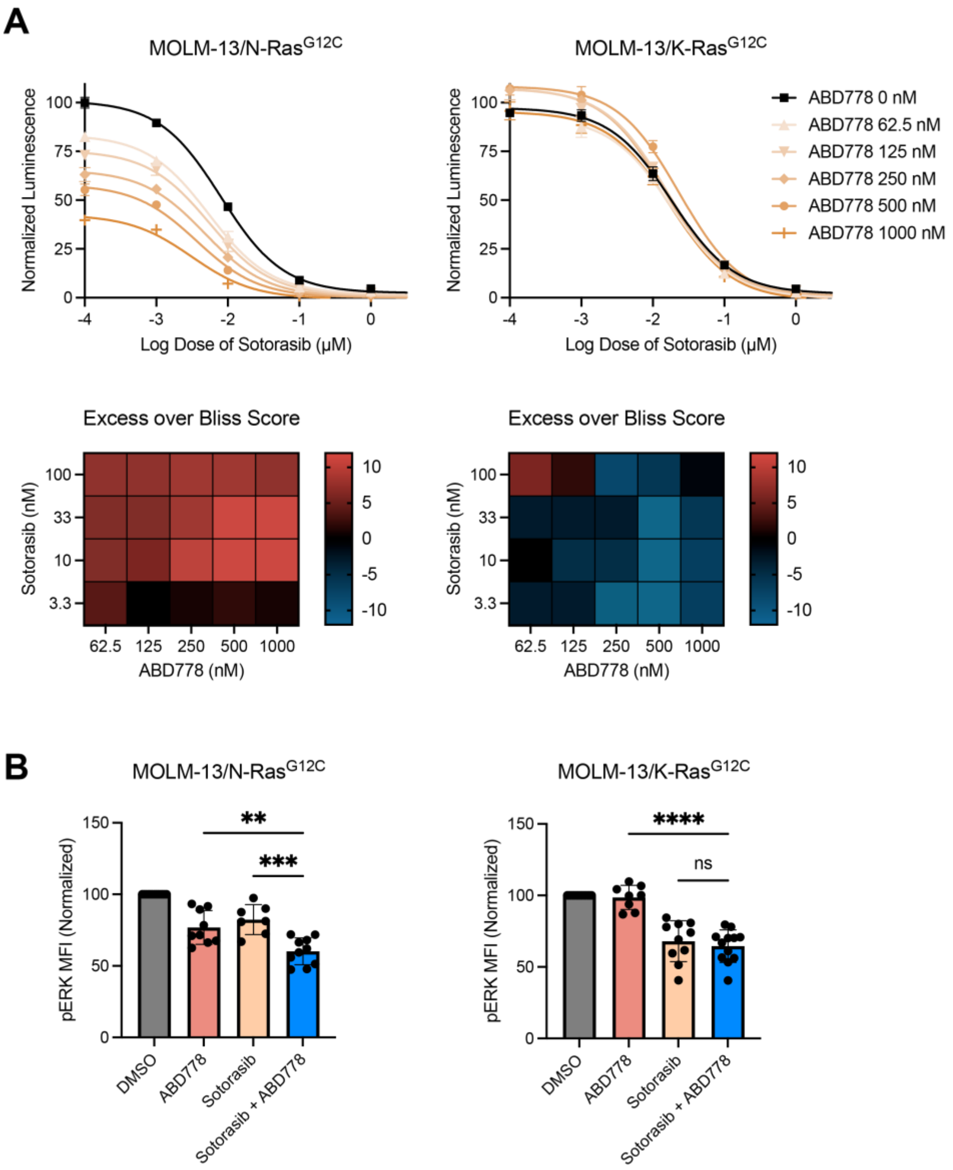


**Figure S14. ABD778 cooperates with sotorasib to repress *NRAS*$^{G12C}$ AML growth and pERK levels. (A)** Bliss independence analysis of MOLM-13/N-Ras$^{G12C}$ (left) and MOLM13/K-Ras$^{G12C}$ (right) cells that were treated with doxycycline (Dox) for 24 h to induce N-Ras$^{G12C}$ or K-Ras$^{G12C}$ expression and then exposed to a range of ABD778 and sotorasib doses. (**B)** pERK levels were measured by phospho-flow cytometry in MOLM13/N-Ras$^{G12C}$ (left) and MOLM13/K-Ras$^{G12C}$ (right) cells that were exposed to the ABD778 and/or sotorasib doses shown for 4 hours (n = 6-14).

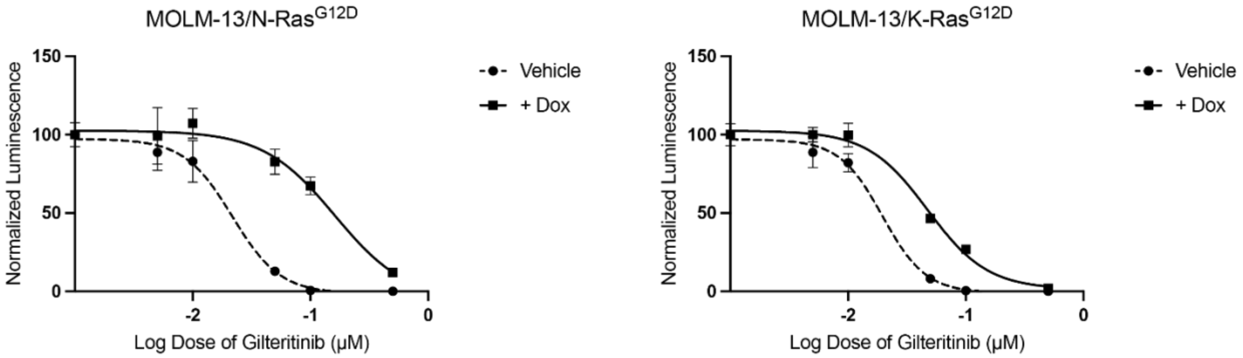


**Figure S15. Sensitivity of parental MOLM-13 cells and of cells engineered to express oncogenic N-Ras$^{G12D}$ or K-Ras$^{G12D}$ to giltiritinib.** MOLM-13 cells were treated with doxycycline (Dox) for 24 hours to induce exogenous N-Ras$^{G12D}$ (left) or K-Ras$^{G12D}$ (right) expression and then exposed a range of gilteritinib doses for 72 hours. Proliferation was assessed by Cell Titer-Glo.

**Table S1**

| Mutation | Cytokine | Drug | HM4483 | HM1989 | HM2491 | Average |
|---|---|---|---|---|---|---|
| *NRAS* | None | DMSO | 6, 5 | 0, 0 | 16, 10 | 6.2 |
| | | ABD778 | 0, 0 | 0, 0 | 7, 6 | 2.2 |
| | GM-CSF | DMSO | 23, 16 | 7, 7 | 57, 36 | 24.3 |
| | | ABD778 | 2, 3 | 0, 0 | 29, 31 | 10.8 |

| Mutation | Cytokine | Drug | HM4208 | HM1959 | HM4210 | Average |
|---|---|---|---|---|---|---|
| *KRAS* | None | DMSO | 48, 64 | 6, 6 | 12, 8 | 24.0 |
| | | ABD778 | 45, 52 | 5, 7 | 5, 8 | 20.3 |
| | GM-CSF | DMSO | 236, 228 | 24, 24 | 41, 28 | 96.8 |
| | | ABD778 | 253, 214 | 29, 23 | 39, 40 | 99.7 |

Legend**.** Number of GFU-GM colonies formed by JMML patient samples harboring somatic *NRAS* (HM4483, HM1989, HM2491) or *KRAS* (HM4208, MH1959, HM4210) mutations in methylcellulose medium, in the presence and absence of GM-CSF.

**Table S2**

| Leukemia | Driver Mutation | Drug and Dose | Vehicle Median Survival | PD901 Median Survival | Difference | P value |
|---|---|---|---|---|---|---|
| 6695 | $Nras^{G12D}$ | PD901 5 mg/kg/day | 10 days | 21 days | 11 days | 0.0025 |
| 6606 | $Nras^{G12D}$ | PD901 5 mg/kg/day | 16 days | 26 days | 10 days | 0.01 |
| 21B | $Kras^{G12D}$ | PD901 5 mg/kg/day | 6 days | 17 days | 11 days | 0.0027 |
| 6768 | $Nras^{G12D}$ | PD901 5 mg/kg/day | 20 days | 23 days | 3 days | 0.2 |
| 63A | $Kras^{G12D}$ | PD901 5 mg/kg/day | 8 days | 19 days | 11 days | 0.0143 |

Legend. Responses of primary mouse AMLs to continuous, single agent treatment with PD901 (5 mg/kg/day) as reported in Burgess MR, et al.. *Blood*. 2014;124(26):3947-3955. and Burgess MR,, et al. *Cell*. 2017;168(5):817-829.e15. P values calculated by log rank (Mantel-Cox) test.

**Table S3**

| Leukemia | Driver Mutation | Drug and Dose | Vehicle Median Survival | PD901 Median Survival | Difference | P value |
|---|---|---|---|---|---|---|
| 6695 | *Nras*$^{G12D}$ | PD901 2.5 mg/kg/day | 11 days | 19 days | 8 days | 0.0027 |
| | | ABD778 60 mg/kg/day | 11 days | 14 days | 3 days | 0.0027 |
| | | PD901 2.5 mg/kg/day + ABD778 60 mg/kg/day | 11 days | 32 days | 21 days | 0.0027 |
| 6768 | *Nras*$^{G12D}$ | PD901 2.5 mg/kg/day | 18 days | 34 days | 16 days | 0.0027 |
| | | ABD778 60 mg/kg/day | 18 days | 24 days | 6 days | 0.0027 |
| | | PD901 2.5 mg/kg/day + ABD778 60 mg/kg/day | 18 days | 38 days | 20 days | 0.0027 |
| 6606 | *Nras*$^{G12D}$ | PD901 2.5 mg/kg/day | 11 days | 19 days | 8 days | 0.0026 |
| | | ABD778 60 mg/kg/day | 11 days | 14 days | 3 days | 0.0026 |
| | | PD901 2.5 mg/kg/day + ABD778 60 mg/kg/day | 11 days | 41 days | 30 days | 0.0026 |
| 21B | *Kras*$^{G12D}$ | PD901 2.5 mg/kg/day | 4 days | 14 days | 10 days | 0.0023 |
| | | ABD778 60 mg/kg/day | 4 days | 5 days | 1 day | 0.5485 |
| | | PD901 2.5 mg/kg/day + ABD778 60 mg/kg/day | 4 days | 10 days | 6 days | 0.0023 |
| 63A | *Kras*$^{G12D}$ | PD901 2.5 mg/kg/day | 9 days | 22 days | 13 days | 0.0101 |
| | | ABD778 60 mg/kg/day | 9 days | 7 days | –2 days | 0.2228 |
| | | PD901 2.5 mg/kg/day + ABD778 60 mg/kg/day | 9 days | 16 days | 7 days | 0.0514 |

Legend. Responses of cohorts of recipient mice (n = 5) that were transplanted with individual primary *Nras*$^{G12D}$ or *Kras*$^{G12D}$ AMLs and assigned to receive control vehicle, ABD778, PD901, or PD901 + ABD778. P values calculated by log rank (Mantel-Cox) test.

**Table S4**

| Leukemia | Driver Mutation | Mouse ID | Treatment | Gene and Mutation | Vehicle VAF | Drug Treated VAF |
|---|---|---|---|---|---|---|
| 6606 | *Nras*$^{G12D}$ | 13 | PD901 + ABD778 | *Nras*$^{G12D}$ | 0.61 | 0.90 |
| | | | | *Asxl1*$^{I594fs}$ | 0.48 | 0.48 |
| | | | | *Kras*$^{A146T}$ | ND | 0.06 |
| | | 16 | PD901 | *Nras*$^{G12D}$ | 0.61 | 0.85 |
| | | | | *Asxl1*$^{I594fs}$ | 0.48 | 0.42 |
| | | 18 | PD901 + ABD778 | *Nras*$^{G12D}$ | 0.61 | 0.94 |
| | | | | *Asxl1*$^{I594fs}$ | 0.48 | 0.48 |
| | | | | *Map2k1*$^{50\_63del}$ | ND | 0.07 |
| | | | | *Braf*$^{G503E}$ | ND | 0.18 |
| | | 20 | PD901 + ABD778 | *Nras*$^{G12D}$ | 0.61 | 0.79 |
| | | | | *Asxl1*$^{I594fs}$ | 0.48 | 0.44 |
| 6768 | *Nras*$^{G12D}$ | 13 | PD901 + ABD778 | *Nras*$^{G12D}$ | 0.40 | 0.55 |
| | | | | *Cebbpa*$^{A83T}$ | 0.45 | 0.50 |
| | | 15 | PD901 + ABD778 | *Nras*$^{G12D}$ | 0.40 | 0.52 |
| | | | | *Cebbpa*$^{A83T}$ | 0.45 | 0.42 |
| | | 18 | PD901 + ABD778 | *Nras*$^{G12D}$ | 0.40 | 0.49 |
| | | | | *Cebbpa*$^{A83T}$ | 0.45 | 0.48 |
| | | | | *Braf*$^{E421V}$ | ND | 0.03 |
| 6695 | *Nras*$^{G12D}$ | 13 | PD901 + ABD778 | *Nras*$^{G12D}$ | 1.00 | 1.00 |
| | | | | *Flt3D*$^{838E}$ | ND | 0.03 |
| | | | | *Fgfr1*$^{N506H}$ | ND | 0.29 |
| | | 15 | PD901 + ABD778 | *Nras*$^{G12D}$ | 1.00 | 0.99 |
| | | | | *Pik3ca*$^{F977Y}$ | ND | 0.04 |

Legend. Mutations were called from exome sequencing of genomic DNA that was isolated from bone marrow cells collected at euthanasia from mice treated with vehicle, PD901 or the PD901 + ABD778 combination. ND = not detected.

### Supplemental Methods

### Synthesis and Characterization of ABD778

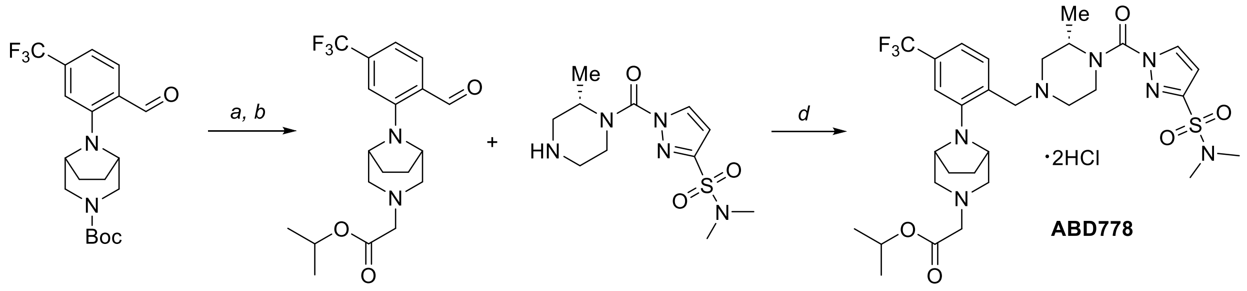


*Step A: 2-((1R,5S)-3,8-diazabicyclo[3.2.1]octan-8-yl)-4-(trifluoromethyl)benzaldehyde hydrochloride*

To a solution of *tert*-butyl 8-[2-formyl-5-(trifluoromethyl)phenyl]-3,8-diazabicyclo[3.2.1]octane-3-carboxylate (75.0 g, 195 mmol, 1.00 equiv) in DCM (200 mL) was added dropwise HCl in 1,4-dioxane (375 mL, 4 N) at 0 °C. The reaction was stirred at room temperature for 12 h before concentrating under reduced pressure to give 2-((1R,5S)-3,8-diazabicyclo[3.2.1]octan-8-yl)-4-(trifluoromethyl)benzaldehyde hydrochloride (64.0 g, crude) as a yellow solid. LCMS (ESI, *m/z*): 285 $[M+H]^+$.

*Step B: isopropyl 2-{8-[2-formyl-5-(trifluoromethyl)phenyl]-3,8-diazabicyclo[3.2.1]octan-3-yl}acetate*

A mixture of 2-((1R,5S)-3,8-diazabicyclo[3.2.1]octan-8-yl)-4-(trifluoromethyl)benzaldehyde hydrochloride (64.0 g, 225 mmol, 1.00 equiv) , isopropyl 2-bromoacetate (48.9 g, 270 mmol, 1.20 equiv) and $K_2CO_3$ (93.2 g, 675 mmol, 3.00 equiv) in DMF (1000 mL) was stirred at 100 °C for 12 h. The reaction was quenched with water (500 mL). The resulting solution was extracted with EtOAc (3 x 500 mL) and the combined organic layers were washed with brine (2 x 200 mL), dried over anhydrous $Na_2SO_4$ and concentrated under reduced pressure. The residue was chromatographed on a silica gel column with petroleum ether:EtOAc (8:1) to give isopropyl 2-{8-[2-formyl-5-(trifluoromethyl)phenyl]-3,8-diazabicyclo[3.2.1]octan-3-yl}acetate (65.8 g, 76% yield) as a yellow solid. LCMS (ESI, *m/z*): 385 $[M+H]^+$.

*Step C: isopropyl 2-[8-(2-{[(3S)-4-[3-(dimethylsulfamoyl)pyrazole-1-carbonyl]-3-methylpiperazin-1-yl]methyl}-5-(trifluoromethyl)phenyl)-3,8-diazabicyclo[3.2.1]octan-3-yl]acetate dihydrochloride (**ABD778**)*

To a stirred mixture of isopropyl 2-{8-[2-formyl-5-(trifluoromethyl)phenyl]-3,8-diazabicyclo[3.2.1]octan-3-yl}acetate (35.0 g, 91.0 mmol, 1.00 equiv) and (S)-N,N-dimethyl-1-(2-methylpiperazine-1-carbonyl)-1H-pyrazole-3-sulfonamide hydrochloride[1] (30.2 g, 100 mmol, 1.10 equiv) in MeOH (700 mL), was added AcOH (4 mL) at room temperature. The reaction was stirred for 12 h at room temperature prior to addition of $NaBH_3CN$ (17.2 g, 273 mmol, 3.00 equiv) at room temperature. The reaction was stirred at room temperature for 12 h before concentrating under reduced pressure. The residue was purified by reverse flash chromatography with the following conditions: column, C18 column; mobile phase, MeCN in water, 60% to 70% gradient in 10 min; detector, UV 220 nm. The purifications were collected and lyophilized. The residue was dissolved into $Et_2O$ (500 mL) and treated with HCl (17.9 mL, 2 N in $Et_2O$, 2 equiv). The resulting mixture was stirred for 2 h and concentrated to provide isopropyl 2-[8-(2-{[(3S)-4-[3-(dimethylsulfamoyl)pyrazole-1-carbonyl]-3-methylpiperazin-1-yl]methyl}-5-(trifluoromethyl)phenyl)-3,8-diazabicyclo[3.2.1]octan-3-yl]acetate dihydrochloride (ABD778) (27.3 g, 40.4% yield) as a white solid.

1H NMR (400 MHz, DMSO-$d_6$) $\delta$ 11.47 (s, 1H), 10.70 (s, 1H), 8.70 – 8.11 (m, 2H), 7.53 (d, $J$ = 8.5 Hz, 1H), 7.23 (s, 1H), 6.91 (d, $J$ = 2.7 Hz, 1H), 5.19 – 4.45 (m, 6H), 4.35 (s, 3H), 4.12 (s, 2H), 3.87 (s, 3H), 3.72 - 3.57 (m, 3H), 3.04 (s, 1H), 2.74 (s, 6H), 2.41 – 2.03 (m, 3H), 1.94 (d, $J$ = 42.3 Hz, 1H), 1.57 (s, 3H), 1.25 (d, $J$ = 6.2 Hz, 6H); 13C NMR (101, MHz, DMSO-$d_6$) $\delta$ 165.56, 150.42, 149.50, 135.14, 133.27, 130.96, 130.63, 128.29, 125.58, 122.87, 120.32, 120.16, 115.37, 108.70, 70.13, 60.64, 58.63, 57.85, 55.37, 54.61, 53.43, 52.06, 38.06, 26.04, 25.81, 21.90, 16.61; HRMS (m/z) [M+H]+ calcd. for $C_{30}H_{43}N_7O_5F_3S$, 670.2998; found 670.2995.

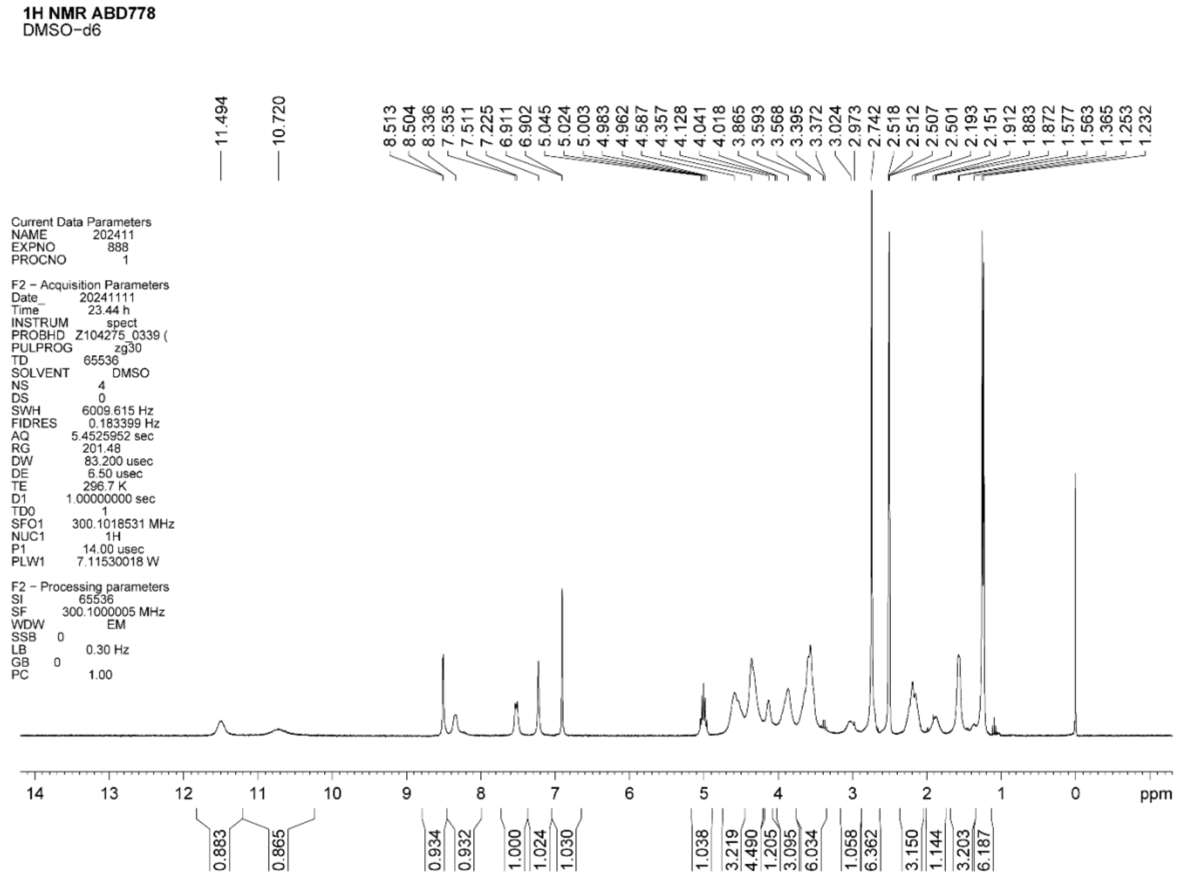

**13C NMR ABD778**
DMSO-d6

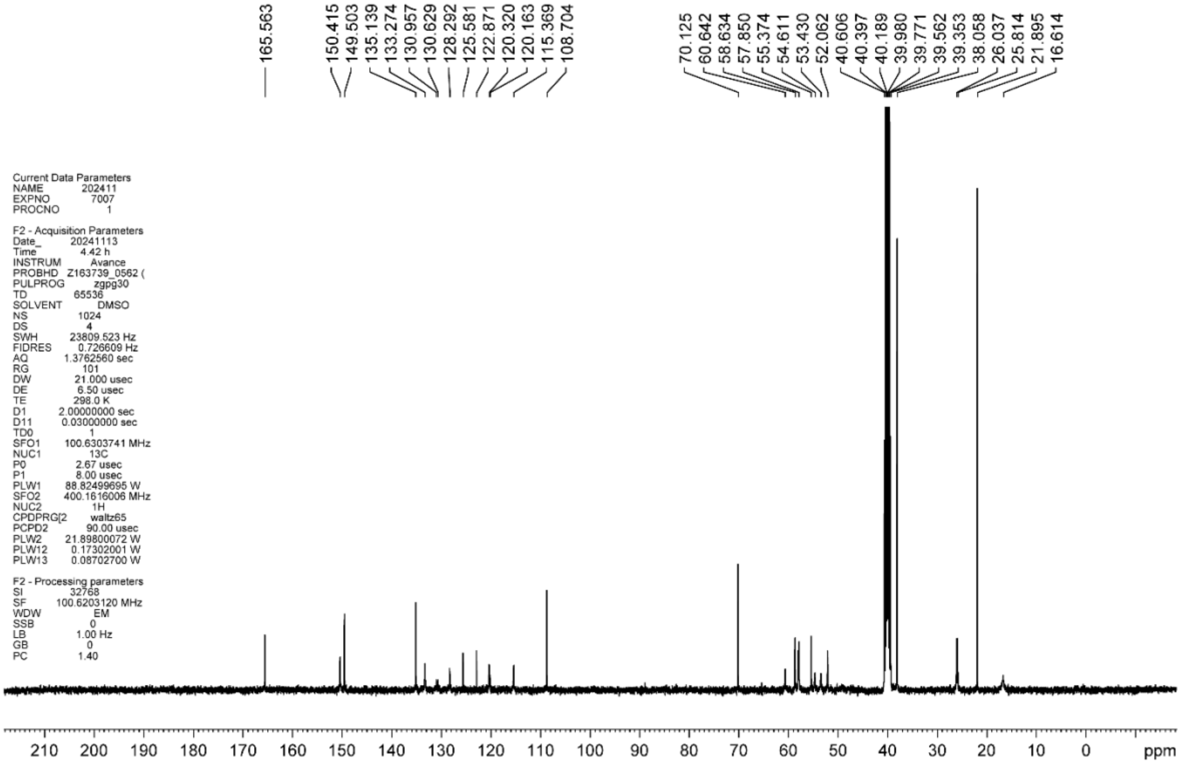
